# The evolution and phylodynamics of serotype A and SAT2 foot-and-mouth disease viruses in endemic regions of Africa

**DOI:** 10.1101/572198

**Authors:** S. Lycett, V.N. Tanya, M. Hall, D. King, S. Mazeri, V. Mioulet, N. Knowles, J. Wadsworth, K. Bachanek-Bankowska, N. Ngwa Victor, K.L. Morgan, B.M.deC. Bronsvoort

**Author notes:** these authors contributed equally to this work (S. Lycett, K.L. Morgan, B.M.deC. Bronsvoort).

## Abstract

Foot-and-mouth disease (FMD) is a major livestock disease with direct clinical impacts as well as indirect trade implications. Control through vaccination and stamping-out has successfully reduced or eradicated the disease from Europe and large parts of South America. However, sub-Saharan Africa remains endemically affected with 5/7 serotypes currently known to be circulating across the continent. This has significant implications both locally for livestock production and poverty reduction but also globally as it represents a major reservoir of viruses, which could spark new epidemics in disease free countries or vaccination zones. This paper describes the phylodynamics of serotypes A and SAT2 in Africa including recent isolates from Cameroon in Central Africa. We estimated the most recent common ancestor for serotype A was an East African virus from the 1930s compared to SAT2 which has a much older common ancestor from the early 1700s. Detailed analysis of the different clades shows clearly that different clades are evolving and diffusing across the landscape at different rates with both serotypes having a particularly recent clade that is evolving and spreading more rapidly than other clades within their serotype. However, the lack of detailed sequence data available for Africa seriously limits our understanding of FMD epidemiology across the continent. A comprehensive view of the evolutionary history and dynamics of FMD viruses is essential to understand many basic epidemiological aspects of FMD in Africa such as the scale of persistence and the role of wildlife and thus the opportunities and scale at which vaccination and other controls could be applied. Finally we ask endemic countries to join the OIE/FAO supported regional networks and take advantage of new cheap technologies being rolled out to collect isolates and submit them to the World Reference Laboratory.

## Introduction

Foot-and-mouth disease (FMD) is a one of the most important livestock diseases globally. It is a highly contagious viral disease affecting all cloven hoofed animals, causing direct production losses in adult animals and mortality in young animals, as well as economic losses through trade interruption.^1,2^ It is caused by the foot-and-mouth disease virus (FMDV), a positive-sense single-stranded RNA virus in the genus *Aphthovirus*, family *Picornaviridae.* Though generally considered a single disease because of the common clinical signs,^3^ from an epidemiological and control point of view, it is better considered as at least 7 immunologically distinct diseases^4^ based on the 7 recognised serotypes currently circulating globally.^5^ These are serotypes O and A found around the world, the Southern African Territories (SAT) 1, 2 and 3 serotypes found predominantly in sub-Saharan Africa, serotype C (which has not been reported since 2004^6^) and Asia1 (found in Asia).

FMD has historically been thought of as important only in middle or high income economies with highly improved breeds and intensive production systems. Introductions into these systems can have devastating effects such as in the UK 2001 outbreak.^7^ FMD is often not considered a priority in endemic regions where low production and extensive livestock systems predominate.^8^ However, there is growing evidence to support the alternative argument that at the individual household level FMD is important in terms of direct losses of production,^9^ lower fertility and loss of other livestock services such as draft power.^10^ Furthermore, it has indirect impacts preventing access to international markets, limiting genetic improvement and preventing development of diary production.^2,8,10^ Even in areas such as Kenya with some commercial dairy production, in spite of regular vaccination, outbreaks continue to have significant impacts on production and culling rates.^11,12^ There is now clear economic evidence for low and middle income countries to invest in control by vaccination with a cost benefit ratio of more than 4.^9^ Furthermore, there is a global societal good to control of FMD in endemic areas, as these reservoirs of disease present a clear and present risk to disease free areas or areas using vaccination, through trade in animal products and animal movements as demonstrated by the recent rapid expansion of the serotype O/ME-SA/IND-2001 lineage across the Middle East and North Africa Southeast and East Asia from Indian sub-continent.^13,14^ The best defense for disease free regions is to reduce the risk to themselves through reducing the disease burden in endemic areas.^1^ This needs a strong coordinated international response to first improve our understanding of FMD epidemiology in endemic settings and to then coordinate controls, as was done successfully with rinderpest globally^15^ and FMD in South America, which has reported only a handful of outbreaks in Colombia since 2014.

Sub-Saharan Africa (SSA) is an enormous geographical area with a huge diversity of cultures, cattle rearing practices and wildlife, including the Cape buffalo (an important reservoir of SAT serotypes), which influence the epidemiology of FMD in different regions.^16^ Historically, Africa is thought to be the origin of the SAT viruses, probably from Cape buffalo, which have gradually spread north through East Africa and then into Central and West Africa.^17^ Serotypes O and A are thought to have been introduced to Africa relatively recently and spread south and west through livestock movements.^16,18,19^ Currently 5 of the 7 serotypes of FMD virus (O, A, SAT1, SAT2 and SAT3) are reported to be circulating in SSA^5,16,19^ and West and Central Africa have had at least 4 of these serotypes (O, A, SAT1 and SAT2) in recent years^20–23^ although until recently (2015), SAT1 had not been reported in West Africa since the early 1980s.^16^ However, because much of SSA has endemic FMD there are relatively few submissions to the World Reference Laboratory (WRLFMD), at The Pirbright Institute, from Africa. Therefore, we have very poor understanding of what viruses are circulating at any given time, the flow of the different serotypes and strains across the continent or the scale and means of persistence at a population level. This lack of basic surveillance data on FMD viruses has major implications for control both locally, regionally and internationally.

FMD control has been very successful in many parts of the world, such as Europe and South America and is making significant impacts now in S.E. Asia.^24^ These have largely been based around vaccination campaigns at 4-6 month intervals and movement controls. In addition, recent support from the Food and Agricultural Organization (FAO) through the European Commission for the Control of FMD (EuFMD) together with the World Organization for Animal Health (OIE) has led to the development of the progressive control pathway (PcP) for FMD^24^ that helps countries frame their control efforts and provides realistic step-wise targets for national governments to help focus efforts. A key element in this pathway is the development of robust surveillance systems to allow regular collection of FMD viruses for typing and sequencing as well as vaccination. However, the details of what individual countries in Africa are doing and the number of outbreaks does not appear to be publicly available, although we know the viruses are endemic and that vaccination with a wide range of vaccines is occurring in many countries, even though in a relatively uncoordinated way. This is important both to improve our understanding of the local epidemiology and where national strains fit in the regional picture and also to help improve vaccine selection for regions.^1^ Increasingly, this sequence information is going to be able to be used directly for vaccine selection as predictive models of antigenicity become available,^25,26^ however, to be able to predict the efficacy of vaccines in different settings, a much better understanding of virus strains and their flows across the continent are needed.

The high number of circulating strains in Africa, the presence of wildlife reservoirs and the diversity of antigenic strains has tended to lead to a sense that FMD control in SSA is neither economically worthwhile or logistically feasible. However, through better surveillance information, understanding of livestock movements^27^ and contact with wildlife and international cooperation and coordination it may be possible to isolate and control individual serotypes in certain zones or to use natural geographical barriers to help zone off areas for control. The emerging view is that FMD control produces a significant amount of public good, justifying the need for national and international public investment in control.^2^

In 2012 we conducted a follow-up study to our 2000 study^4,20^ of circulating FMD viruses in the Adamawa and the Northwest Regions of Cameroon, to investigate the genetic changes in the serotypes and strains over the intervening 12 years and understand their relationship to the relatively small number of other sequences submitted from Africa over this period particularly in relation to the North African outbreaks.^28^ This is important as FMD surveillance in SSA is still fragmented and at very low coverage. This paper describes the latest patterns of serotypes A and SAT2 in SSA and estimates the dates of most common ancestors for key lineages and topotypes and discusses the different rates of spread of key lineages and topotypes.

## Results

### Circulation Patterns in Serotype A

A set of 875 serotype A VP1 sequences was compiled from GenBank, the WRLFMD and Cameroon studies, and used to create a maximum likelihood phylogenetic tree (Figure 1) showing the relationship between strains on different continents. The data set consists of African (19%), Asian (63%), European (3%) and South American (15%) sequences, and shows three global clades of predominantly Asian (purple), South American and European (blues) and African (red) taxa. There are three African sequences within the South American and European clade (from Morocco and Libya 1977-1983), and seven African sequences within the Asian clade (from Libya in 2009 and Egypt 2010-2013), showing that there are occasional incursions of non-African lineages into Northern Africa, however the majority of the circulation in Africa is from African strains, and no sequence from elsewhere clusters with the main African clade.

**Figure 1.**
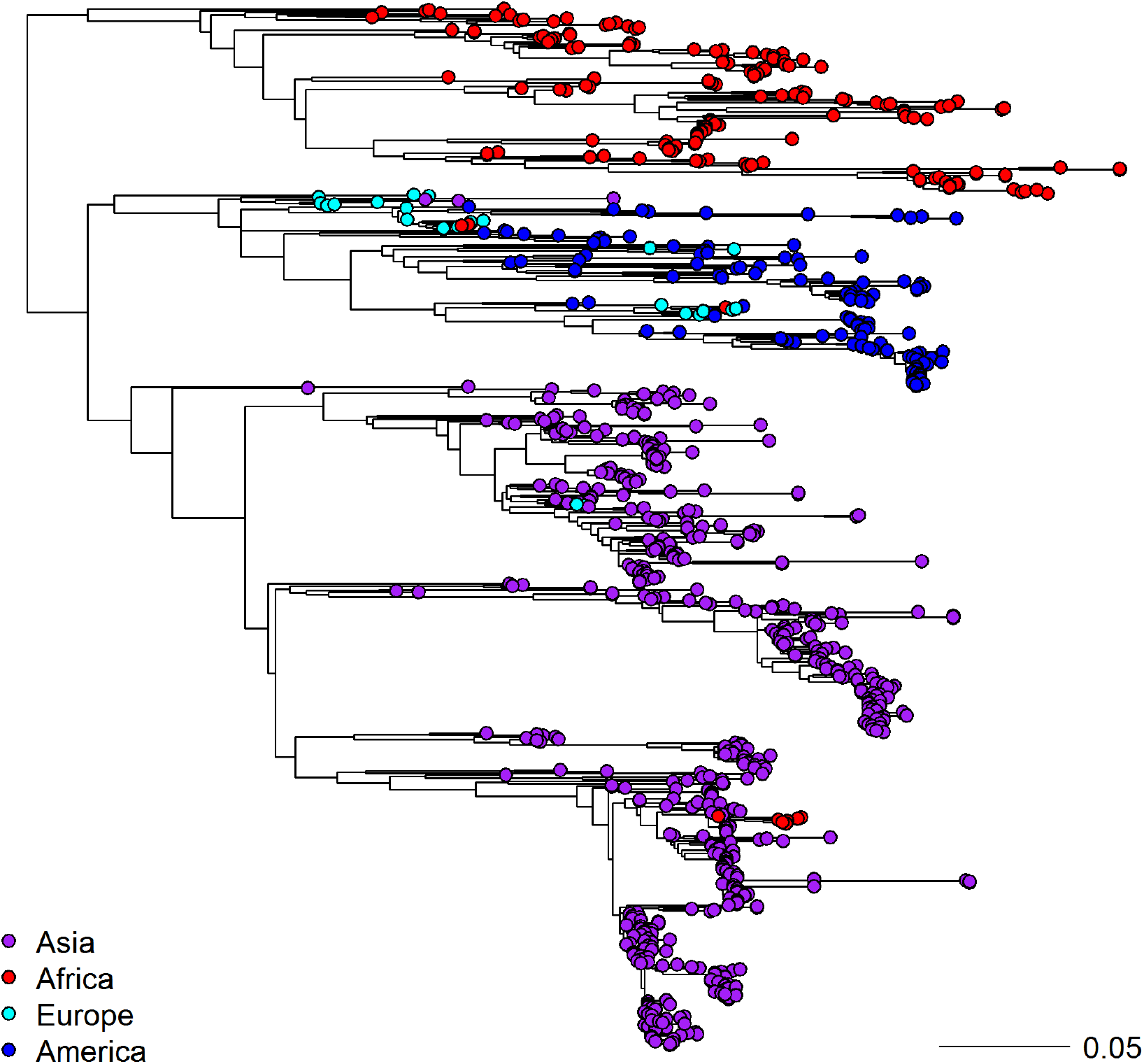
Maximum likelihood phylogenetic tree of worldwide FMD serotype A viruses based on 875 VP1 sequences

Next, to look at the transmission pattern of serotype A within Africa, we used sequences from Africa clustering within the main African clade (154 in total – also see Figure S1) to perform a time resolved phylogeographic analysis using BEAST. Region labels were added to the sequences (Central, Eastern, Northern and Western; there were no Southern), and the number of transmissions between regions were inferred using an asymmetric discrete traits model on the time resolved trees. Figure 2 shows the time resolved tree of the African serotype A sequences, with the branches and nodes coloured by inferred discrete region trait and the size of the ancestral nodes reflecting the probability of the inferred region at that point.

**Figure 2.**
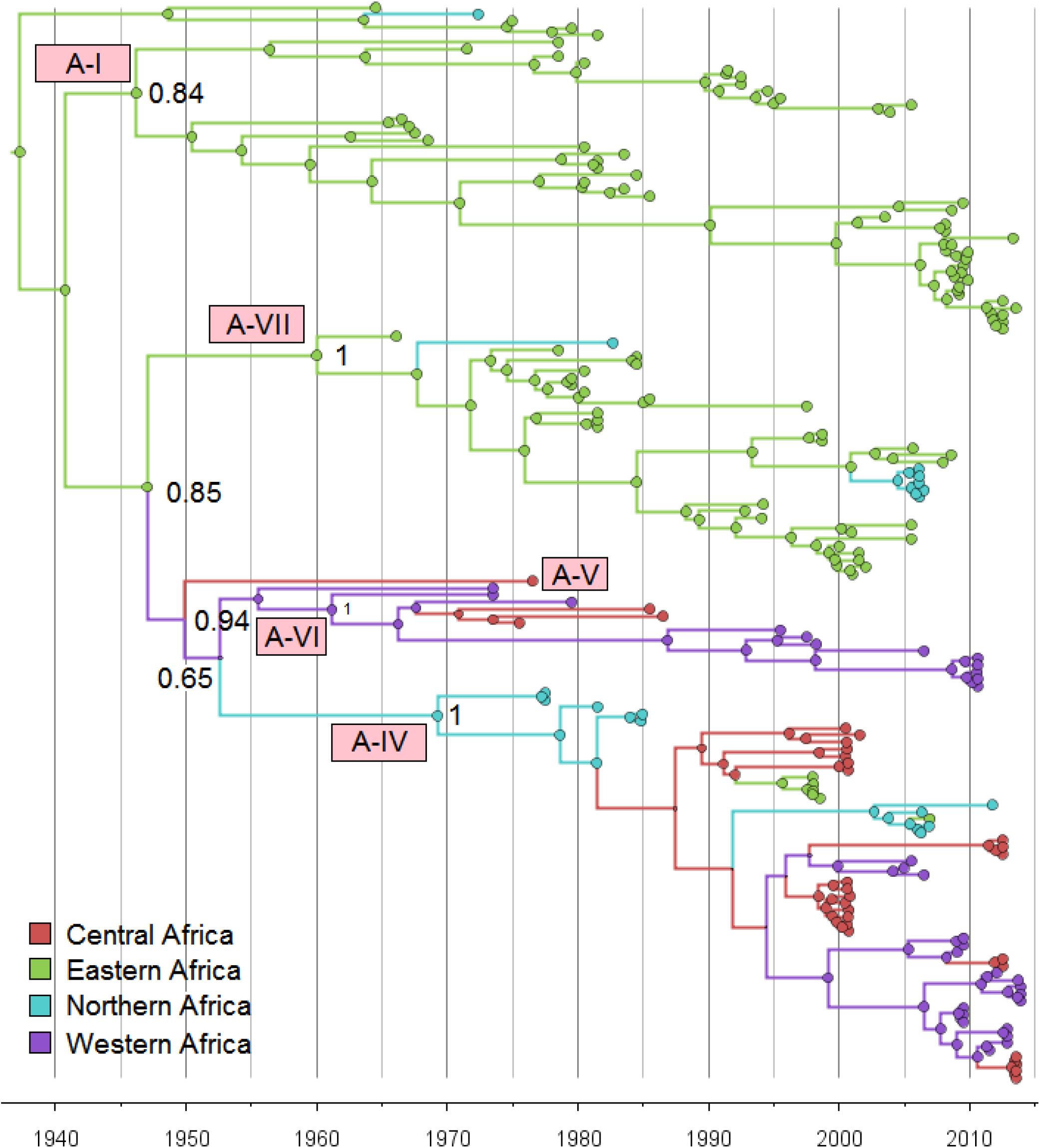
Time scaled tree of 154 serotype A VP1 sequences from Africa

The tree in Figure 2 indicates that all these serotype A sequences have an Eastern Africa most recent common ancestor in the 1930’s (median 1937 with 95% HPD: 1922-1950), and suggests that there is an interchange of viruses between Northern, Central and Western Africa (genotypes A-IV, V and VI), and between Eastern only, or Eastern and Northern Africa separately (genotypes A-I and A-VII).

From the tree, it can also be seen that there is only one case of Central to Eastern transmission (within A-IV) related to a sample from Eritrea in 1998 (A/ERI/3/98), highlighting that although Eastern-Central transmission is possible, it does not seem to occur very frequently.

The lower clade in Figure 2 contains all the Central and Western serotype A strains, and some Northern strains. It is split into two main sub-clades; genotype A-VI contains strains isolated in the 1970s from Cameroon, Nigeria, and Ghana, and also recent viruses isolated from Benin in 2010, whereas A-IV contains strains isolated from Sudan from 1977-1984 and again in 2006, and the majority of the Cameroonian strains from 2000 and 2012-2013.

Figure 3 (and Figure S2) shows the spatial distribution of the sampling locations, together with the approximate spatial extent of the genotypes. From these it can be seen that A-I, II, III and A-VII are in Eastern Africa, and overlap with A-IV in the Northern region only.

**Figure 3.**
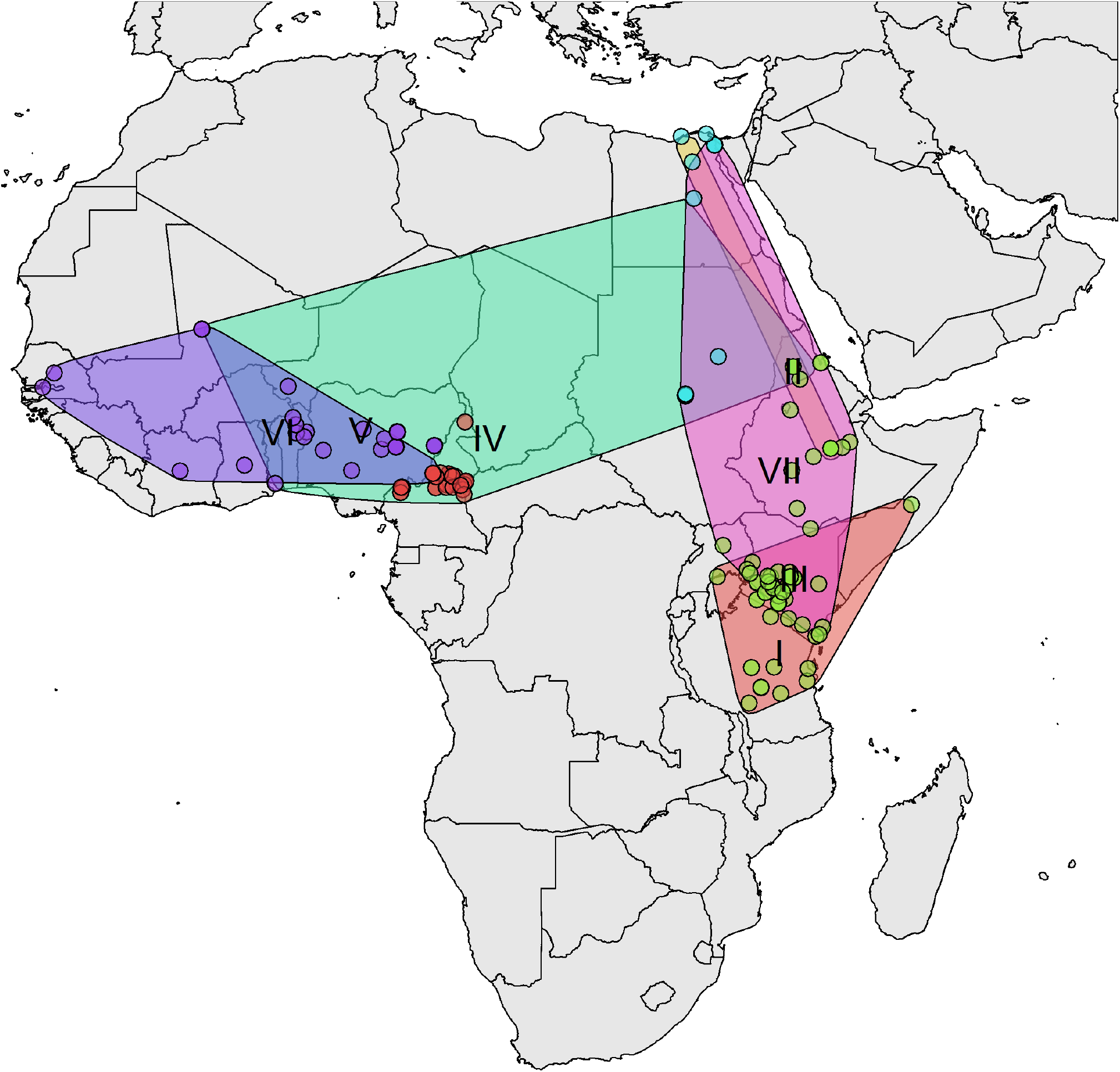
Distribution of A VP1 sequences with approximate spatial extent of the topotypes

Overall, the pattern suggests that the serotype A viruses are potentially circulating in at least 2 quite separate pools in Central-Western Africa and Eastern Africa, and that Cameroon and Central Africa more widely may be an important bridging zone between virus pools (Figure 3). However as with all these analyses in Africa, note the actual virus sampling coverage is extremely low so phylogenetic analyses might give a distorted view of the true transmission pattern.

### Serotype SAT2 Diversity

A set of all available good quality SAT2 VP1 sequences was compiled from GenBank, the WRLFMD and Cameroon studies, of these only 9 were from outside Africa. These were in Saudi Arabia, Bahrain, Yemmen and Palistine (Gaza Strip), showing that the SAT2 serotypes are indeed mostly restricted to Africa with only occasional incursions into the Middle East. Using a set of 334 SAT2 VP1 sequences (see also Figure S3) from Africa only (and removing very similar sequences, see Methods and Figure S4 and S5), Figure 4 shows a maximum likelihood phylogenetic tree with the 14 SAT2 topotypes (clades) indicated,^16^ and Figure 5 (and Figure S6) shows the approximate spatial distribution of these topotype clades from the sampling locations. Similar to the overall pattern observed for serotype A, the SAT2 strains also show distinct topotype clades in Eastern-Southern Africa and Central-Western Africa, and the SAT2 strains from Cameroon fall into topotype VII, which is very widely geographically spread.

**Figure 4.**
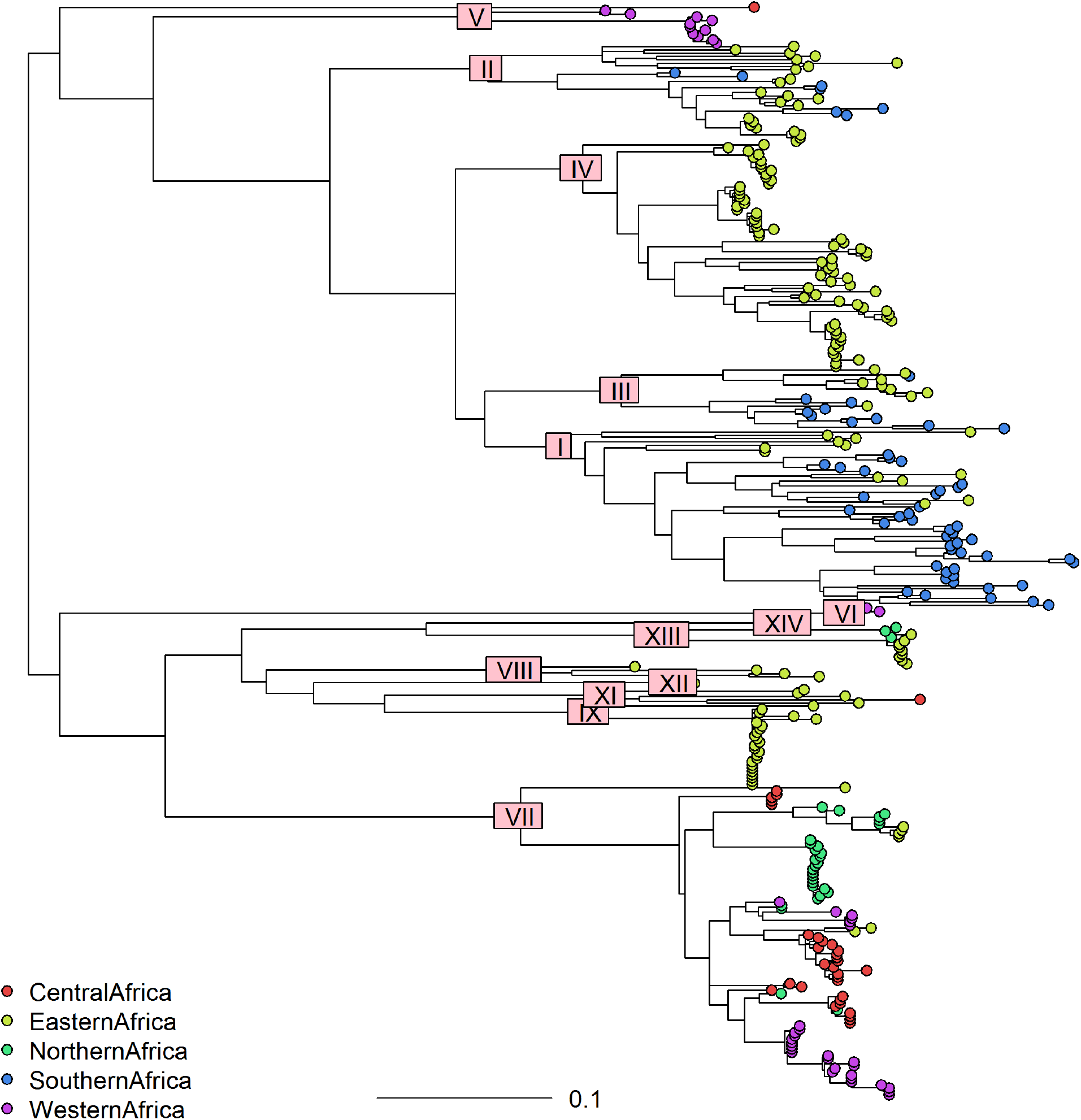
Maximum Likelihood tree of African SAT2 viruses based on 334 VP1 sequences showing topotypes as designated by the FMD World Reference Laboratory

**Figure 5.**
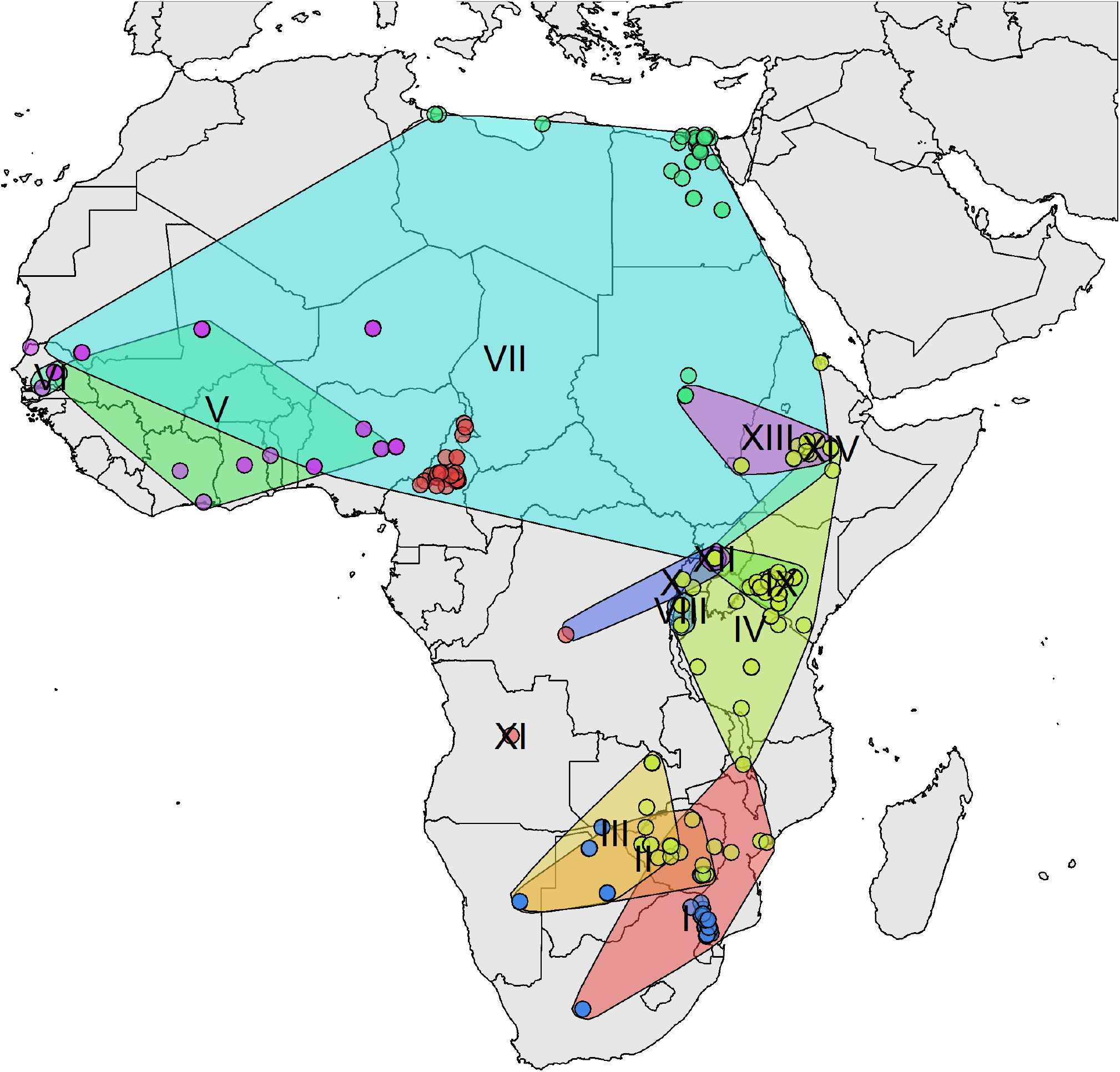
Distribution of SAT2 VP1 sequences with approximate spatial extent of the topotypes

The time resolved maximum clade credibility tree for all 334 SAT2 sequences (Figure S5) additionally indicates that the topotype clades are diverse; in contrast to the serotype A tree, the SAT2 tree shows a much longer time to most recent common ancestor (TMRCA for SAT2 is 1709 with 95% HPD:1502-1814), implying that the SAT2 serotype has been circulating in Africa for at least a few centuries. Topotype VII appears to have a common ancestor from the 1950s (1954: 95% HPD:1906-1984) in East Africa and the more recent isolates in Cameroon from 2012/13 all appear to have a common ancestor with those from 2000 but not apparently directly descended from them.

### Genetic Diversity

The average pairwise distance between all 154 African serotype A amino acids sequences is 13% (28 amino acids – also see Figure S7 and S9), and the average pairwise distance between the 72 sequences in A-IV,V and VI is 8% (20 amino acids). The genetic diversity between SAT2 topotype clades as estimated from amino acid sequences is quite large; and the average pairwise distances between sequences from one topotype to another is 18% (41 amino acids from alignment length of 224), whereas the average pairwise distance within a topotype is 7% (17 amino acids – also see Figure S8 and S9). It is likely that the differences between SAT2 topotypes represent distinct antigenic variants,^26^ and in comparison the average amino acid difference between H3N2 human seasonal influenza vaccine updates is 12 (range 6-20) amino acids in HA1 (length 344, and the most variable part of the HA surface protein, equivalent to the partial VP1 sequences here).

### Diversity and Spatial Diffusion

Figure 6 (and Figure S10) summarise measures of diversity (TMRCA and number of amino acid differences), estimates of the molecular clock evolutionary rate and spatial diffusion rate for topotypes I, IV, VII of SAT2, and A-I, A-VII and A-IV of serotype A using independent BEAST analyses. These clades were screened for recombination, but no recombination was found. The clades contain between 35 (A-VII) and 69 (SAT2-IV) sequences collected over a time span of almost 60 years, however SAT2-I and SAT2-IV have long TMRCAs of 1900 (95% HPD:1830-1943) and 1752 (95% HPD: 1626-1846) respectively, whilst SAT2-VII and A-IV have much more recent TMRCAs of 1977 (95% HPD: 1944-1995) and 1975 (95% HPD: 1973-1978) respectively. The corresponding differences in evolutionary rates are also pronounced – SAT2-VII and A-IV (means of 7e-3 and 9e-3 substitutions/site/year) have much higher rates than SAT2-I and SAT2-IV (means of 3e-3 and 7e-4 respectively). Similarly the spatial diffusion rate estimates of SAT2-VII, A-VII and A-IV are much faster (means of 327, 110 and 209) than SAT2-I, SAT2-IV and A-I (means of 20, 9, and 34). Overall these results point to clades containing central, western and northern African strains, particularly SAT2-VII and A-IV having a faster and wider spatial spread, and faster evolutionary rate than the large eastern and southern SAT2-I, SAT2-IV and A-I clades.

**Figure 6.**
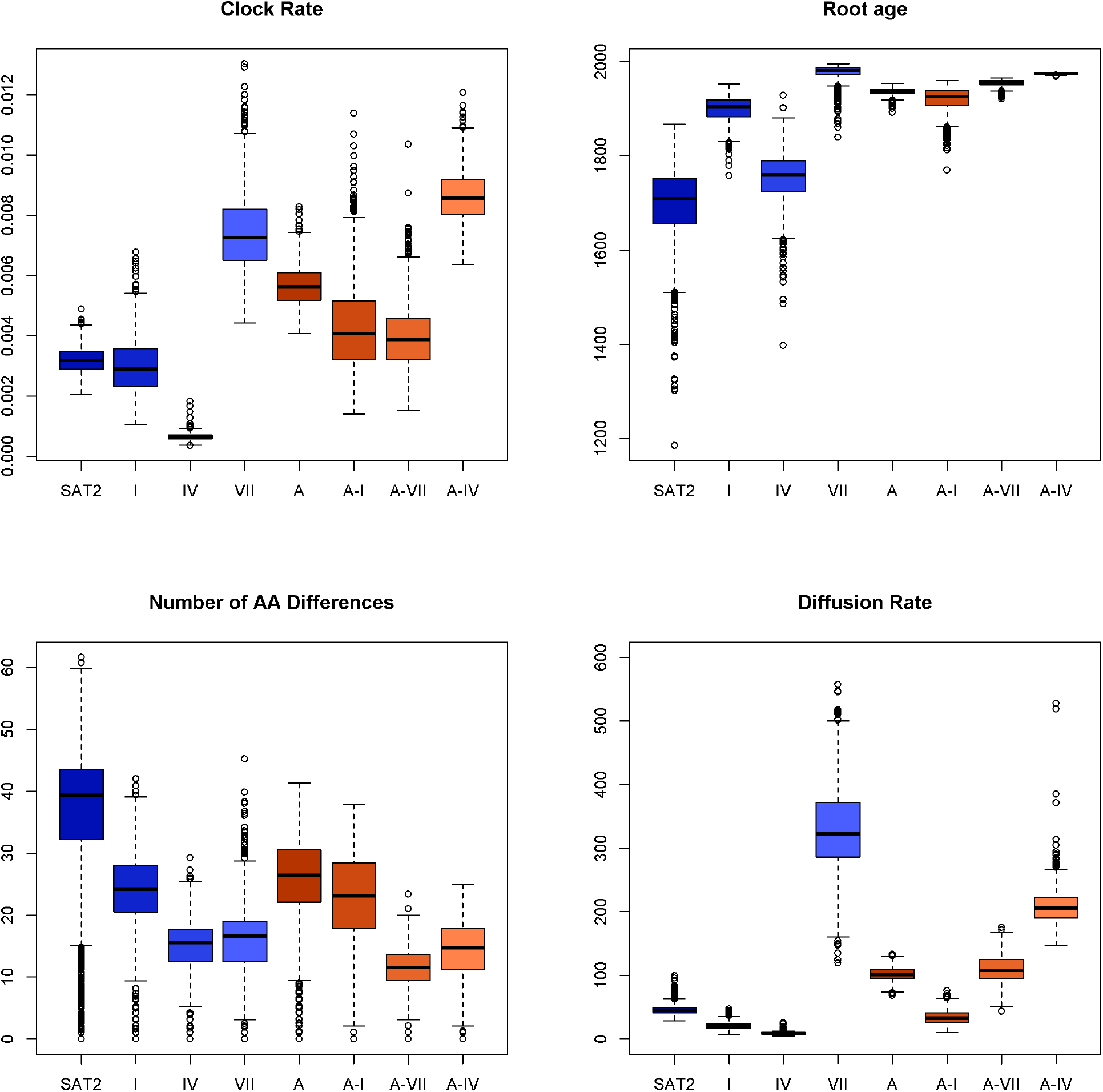
Comparison of SAT2 and A sequences (whole datasets) and individual clades showing overall clock rate, root age, average number of amino acid differences within each clade and rate of spatial diffusion.

### Sites under positive selection

To investigate the selection pressures on the A and SAT2 sequences, measures of non-synonymous to synonymous changes per site were calculated for each clade. Table 1 (and Figure S11) summarises the number of sites under positive (diversifying) selection in the two antigenic regions and the sites under positive and negative (purifying) selection in the receptor binding site RGD, and table 2 summarises the sites under directional selection.

**Table 1.** Sites under selection by clade. Sites are counted as positive (or negative) if any one of the following tests is significant SLAC ≤0.01, FEL ≤0.01, MEME ≤0.01, FUBAR ≥ 0.99 (equivalent to p-value ≤0.05 with Bonferroni correction).

| Clade | #Pos. | Antigenic Region .1 | Antigenic Region .2 | Other | #Neg. | RGD Neg. |
| --- | --- | --- | --- | --- | --- | --- |
| SAT2-I | 1 | - | - | 176 | 152 | 144,145,146 |
| SAT2-IV | 3 | 149 | - | 130,217 | 96 | 145,146 |
| SAT2-VII | 1 | 161 | - | - | 84 | 144,146 |
| A-I | 1 | - | 197 | - | 83 | 146 |
| A-VII | 1 | - | 198 | - | 27 | - |
| A-IV | 2 | - | 196 | 43 | 69 | 145,146 |

**Table 2.** Sites under directional selection by clade. Type of selection is indicated CE: Convergent evolution, SS: Selective Sweeps, FD: Frequency Dependeny selection

| Clade | #Dir | Antigenic Region 1 | Antigenic Region 2 | Other |
| --- | --- | --- | --- | --- |
| SAT2-I | 2 | 142(CE) | - | 86(CE) |
| SAT2-IV | 2 | - | 201(SS),202(SS) | - |
| SAT2-VII | 2 | - | 202(SS) | 56(SS) |
| A-I | 0 | - | - | - |
| A-VII | 1 | 149(CE) | - | - |
| A-IV | 7 | 139(SS),142(SS),149(CE),156(FD) | - | 45(SS),68(SS),171(SS) |

In all clades there were many sites under negative selection as expected, and the RGD site is conserved and under negative selection in the SAT2 serotypes. However, in serotype A not all positions of the RGD site are conserved or are detected as under negative selection, particularly in clade A-VII. None of the clades have positively selected RGD sites. In clades SAT2-IV, SAT2-VII, and in the A clades, there are a few sites under positive selection and these tend to occur in the two surface exposed antigenic regions identified by Reeve et al.^25,26^ Additionally, several sites identified as under directional selection also occur in these antigenic regions. Interestingly, A-IV has more sites under directional selection than the other clades, including sites near the RGD sites. The sites under directional selection in the antigenic regions suggest that particular antigenic variants might be becoming more widespread over time (especially for A-IV).

## Discussion

FMD remains a major global problem both for developed economies, where it has largely been eradicated but remain under threat of reintroductions, and in low and middle income economies where it is under varying levels of control but still impacts production and causes trade blocks. SSA remains as a major reservoir of at least 5 serotypes of FMDVs and until controls are improved in these areas it will remain a threat to global livestock production. This analysis using phylogeographical approaches highlights a number of key issues in FMDV epidemiology in SSA. It suggests that serotype A may have only been introduced relatively recently, and that East Africa may have been the origin, in the late 1930s, for the North, Central and West African isolates studied. In contrast the SAT2 serotype appears to have much older origins going back hundreds of years (and may have originated in Africa thousands of years ago in wildlife). Interestingly, the phylogenetic tree is also dominated by East African viruses and suggests that the appearance of SAT2 in Central Africa may be linked to East Africa and a common ancestor from the 1970s. The isolates from 2005 and later in Central and West Africa and the outbreaks in North Africa all appear to have a common ancestor in the 1970s or 1980s.

It is impossible to fully understand the degree of virus exchange between different regions as these areas are considered endemic for FMD and have not been routinely submitting isolates or reporting outbreaks. However, it is clear that from time to time there are transmission events between regions. However, again the lack of available isolates makes interpretation of these data difficult and there is a real risk of over interpretation. Furthermore, it is also not possible to say if vaccination is driving evolution of viruses in the region due to lack of data on outbreaks, vaccination coverage and vaccine use.

This analysis also highlights that although there is cross over between regions there appear to be antigenically distinct viruses circulating in different parts of Africa and therefore there needs to be more work done to identify strains that will provide protection in these different regions. Additionally, it seems that the clades SAT2-VII and A-IV containing Central, Western and Northern strains are not only distinct from their Eastern and Southern counter parts, but appear to be undergoing faster evolution and faster spatial spread. There are signatures of positive and directional selection in antigenic regions close to the receptor binding site of the VP1 protein in these clades, which are consistent with the faster evolutionary rates, and consequently provide an intriguing hypothesis to be tested in the future: that the increased spread of these clades could be due to a genetic change in the virus. An alternative hypothesis is that the increased spread of these clades is due to increased cattle movements (or human movements of animal products) particularly between the Central and Northern African regions.

This analysis has highlighted our lack of data and understanding of the epidemiology of FMD viruses in SSA and points to the need for a major scaling up of surveillance and sampling across SSA to start to understand key questions on the flow and circulation of FMD viruses in SSA and the scale of persistence of strains and the role of wildlife. Without much higher coverage and spatial resolution of phylogenetic data it will be difficult to design strategies for FMD control across the continent. However, the development of simple to use tools such as the lateral flow device^29^ which could be extended to include African serotype specific bands^30^ would give local veterinary services the means to type viruses in the field. This would be a major step forward for local veterinary staff and livestock keepers to understand the complexity of the epidemiology of FMD due to the number of different serotypes and start the education process needed to drive controls from the bottom up. Furthermore these devices can be safely and cheaply submitted to reference laboratories where the virus can be eluted and sequenced.^31^ Additionally mobile field sequencing technology is also now available as has been used in the Ebola outbreak in West Africa.^32^ This also has the potential to increase the number of sequences, particularly whole genomes and thus help improve local understanding as well as international understanding of the phylogeography of FMD viruses, recombination events and vaccine driven selection in SSA.

## Methods

Most of the sequences used in this study were taken from existing databases but a small set of new isolates from Cameroon were collected for this study. Details of field sample collection and processing are given in the supplementary material. This field work was carried out with the Cameroonian Ministere de l’Elevage des Peches et Industries Animales (MINEPIA) staff, following local guidelines and regulations, as part of their disease surveillance activities and had approval from the Cameroonian Academy of Science.

### GenBank Data

Global data sets of the VP1 region from serotypes A and SAT2 were downloaded from GenBank; an initial search for VP1 (or 1D) regions gave >900 and >400 sequences respectively. Of these we included good quality sequences in the final set with at least 240 nucleotides and both date and country information, together with the new sequences from Cameroon, resulting in data sets of 875 A and 394 SAT2 VP1 sequences (95% and 75% of which had at least 600 nucleotides respectively).

African sequences were extracted from the global data sets and used for more detailed analysis. For the serotype A sequences, the 10 sequences from Northern Africa (6 Egypt, 2 Libya, 2 Morocco) which clustered with non-African sequences were excluded, as was Kenya/1965 (the outgroup). For the SAT2 sequences, the 9 of non-African origin were excluded (all from the Middle East). 9 serotype A and 9 SAT2 sequences from previously unreported isolates are also included in the data sets. These were collected in 2012 from the Adamawa and North West Regions of Cameroon and the sample codes and accession numbers of the sequenced samples are given in Table 3. A further 3 sequences from the Extreme North Region of Cameroon were provided by Ludi et al.^33^ Sequences from the same location, isolated within one week of each other and with less than 3 nucleotides difference were considered the same and the duplicates were removed from the final data set, leaving 154 serotype A sequences in the main monophyletic African clade, and 334 serotype SAT2 sequences.

**Table 3.**
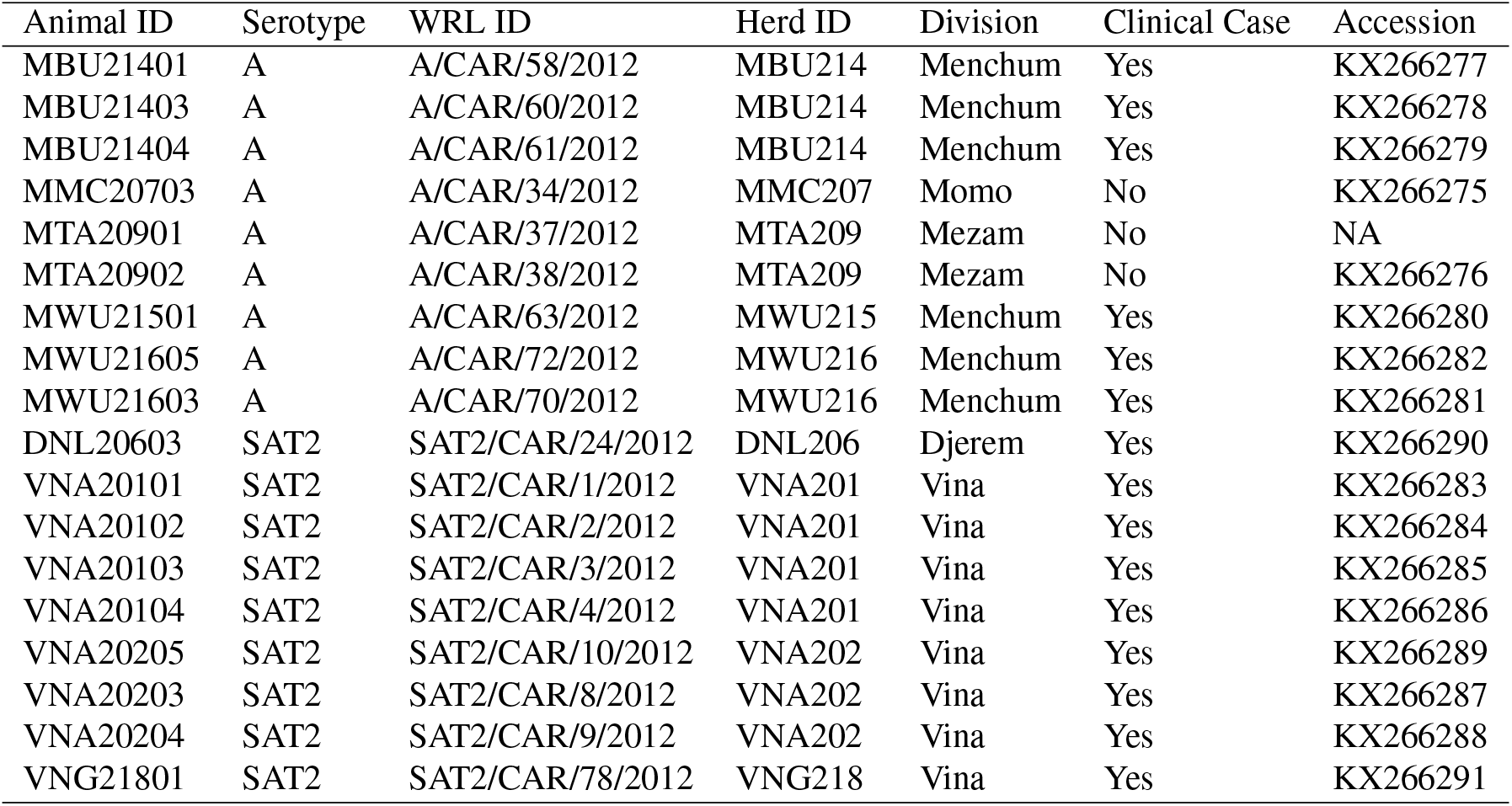
Sample and herd codes for FMD isolates generated from 2012 sampling in Cameroon

### Phylogenetic analysis

#### Alignment

Initial nucleotide alignment was performed using ClustalW in BioEdit, the sequences were manually adjusted to the coding frame, and then aligned using MUSCLE in MEGA6 on translated sequences.

#### Maximum Likelihood Trees

Maximum Likelihood trees on the global data sets were generated using RAxML with the general time reversible (GTR) model and gamma distributed site to site rate variation (4 categories) and 100 bootsraps.

#### Phylodynamics and Phylogeography

Time scaled phylogenetic trees were inferring using BEAST 1.8.^34^ Initial model selection was performed by comparing the fit of substitution models, clock models and effective population size tree prior models calculated using AICM with 100 bootstraps in Tracer 1.6, and Path sampling (PS) and Stepping Stone sampling (SS) in BEAST 1.8.^35,36^ Models were fitted on serotype A sequences using an MCMC chain with length 50,000,000 sampling every 5,000 and discarding the first 10% of samples as burn-in. The SRD06 substitution model^37^ (which uses two HKY models with site to site rate variation, one model for positions 1 and 2, and another for position 3) was favoured above the GTR model with gamma distributed site to site rate variation (4 categories) with a Bayes Factor > 100 (AICM, PS, SS), and the uncorrelated relaxed lognormal clock model was favoured over the strict clock with Bayes Factor >100 (AICM, PS, SS). The constant population size tree priors were favoured over flexible skygrid effective population size tree priors with Bayes Factor >6 for AICM,^38^ but the reverse was true when using PS and SS (Bayes Factor >10). Therefore following Hall *et al*.^17^ we chose to use the flexible skygrid effective population size over time model for serotype A, although this made less than 1% difference to the TMRCAs for recent clades. Since the SAT2 sequences were very diverse and the TMRCA was much longer than for the serotype A sequences, we used a constant population size model when analysing all the topotypes together but a skygrid model when analysing individual topotypes.

A posterior set of 1000 trees for each of serotype A and SAT2 data sets was composed from a minimum of two independent runs, and the location states were reconstructed upon these trees in BEAST 1.8 using (i) asymmetric discrete location trait models with Markov Jumps and (ii) continuous latitude and longitude homogeneous Brownian motion diffusion models^39,40^ (a random jitter of 0.01 was applied to the GPS sampling coordinates in order to give each isolate a unique location).

#### Selection tests

Sites under positive (diversifying) and negative (purifying) selection were detected within the SAT2-I, IV, VII and A-I, IV, VII clades using HyPhy on the DataMonkey server at https://www.datamonkey.org/ to estimate the non-synonymous to synomyous (dN/dS or dN-dS) ratios. SLAC (Single Ancestor Counting), FEL (Fixed Effects Likelihood), MEME (Mixed EffectsModel of Evolution) and FUBAR (Fast Unbiased Bayesian AppRoximation) methods were used (ref1) and a GTR (General Time Reversible, also known as “REV”) model was used as the basis for the codon models. On the HyPhy server, neighbour joining trees are automatically created from the uploaded sequences, and it is these which are used as the phylogeny within the codon-based selection tests. These selection tests have different detailed assumptions and performance depending on the size and variability of the data, but here a site was reported to be under positive or negative selection in the main text if it was significant under any of the tests with a p-value of 0.01 or FUBAR ≥0.99 posterior probability.

The SLAC^41^ method reconstructs the maximum likelihood ancestral sequences at each site in the alignment up the phylogenetic tree according to an overall fitted codon model (disallowing stop codons), and then counts the number of synonymous vs non-synonymous substitutions. This is the fastest and crudest method. FEL^41^ is also a site-by-site model, but here the rates of non-synonymous and synonymous substitution for each site are estimated independently as extra parameters besides the branch lengths and overall site variability. A likelihood ratio test is performed for each site to test whether a one parameter model (non-synonymous rate is the same as the synonymous rate) or a two parameter model (the rates are different) is applicable and hence the type of selection the site is under. In FUBAR,^42^ independent non-synonymous and synonymous substitution rates are also inferred for each site (using Bayesian inference), and these are considered as random effects drawn from overall discretised distributions. MEME^43^ is another random effects model for individual sites, but rather than assuming that the selection pressure must be constant for a site over all time (and all branches), it allows the detection of both pervasive and episodic diversifying evolution along only some of the branches.

To further investigate the type of selection that individual sites were under, we also performed a Directional Evolution for Protein Sequences (DEPS) analysis^44^ on the HyPhy server; this analysis used only amino acid sequences and uploaded rooted neighbour joining trees, created from the nucleotide sequences and the oldest sequence per serotype as outgroup. The best fitting amino acid substitution model was determined from each clade separately and was JTT^45^ for serotype A A-I, IV, VII and SAT2-VII, and WAG^46^ for SAT2-I and IV. DEPS analyses considers the number and type of amino acid transitions per site and reports sites where the transitions indicate a: selective sweep site, consensus replacement site, repeated substitutions site (convergent evolution), rare residue substitution, or highly polymorphic site.

#### Recombination analyses

Since the presence of recombinant sequences can affect the phylogenetic results and results from selection tests, the VP1 sequences from the SAT2-I, IV, VII and A-I, IV, VII clades were screened for recombination using Single Break Point analysis with a GTR model on the HyPhy server, but no recombination was found.

## Acknowledgements

The authors would like to thank the staff of MINEPIA Cameroon who helped identify infected herds to examine and to the the herdsmen for allowing us access to sample their animals. The Roslin Institute receives Institute Strategic Grant funding from the Biotechnology and Biological Sciences Research Council (BBSRC), BBS/E/D/20002173. SL is supported by a Chancellor’s Fellowship from the University of Edinburgh. Sequencing work at the WRLFMD was supported by the Department of Environment, Food and Rural affairs (Project SE2943) and funding provided to the European Commission for Control of FMD (EuFMD) from the European Union. The views herein can in no way be taken to reflect the official opinion of the European Union.

## Author contributions statement

BB, VT and KM conceived the study; SM, VN, VT, KM and BB collected the samples in the field; DPK, VM, NJK, JW and KBB carried out the virology, the molecular analysis and preliminary phylogenetic analysis; SL and MH conducted phylogenetic and BEAST analyses; BB, SL and KM wrote the paper. All authors reviewed the manuscript.

## Additional information

The authors confirm that there are no competing interests that would affect this research work.

## References

1. Kitching, P. et al. Global FMD control-Is it an option? Vaccine 25, 5660–5664 (2007).

2. Knight-jones, T. J. D. & Rushton, J. The economic impacts of foot and mouth disease-What are they, how big are they and where do they occur? Preventive Veterinary Medicine 112, 161–173 (2013). URL http://dx.doi.org/10.1016/j.prevetmed.2013.07.013.

3. Kitching, R. P. & Alexandersen, S. Clinical variation in foot and mouth disease: pigs. Revue Scientifique et Technique de l’Office International des Epizooties 21, 531–538 (2002).

4. Bronsvoort, B. et al. A serological survey for foot-and-mouth disease in wildlife in East Africa. In 2006 Session of the Research Group of the Standing Technical Committee of EUFMD (Paphos, Cyprus, 2006).

5. Rweyemamu, M. et al. Planning for the Progressive Control of Foot-and-Mouth Disease Worldwide. Transboundary and Emerging Diseases 55, 73–87 (2008).

6. Sangula, A. K. et al. Low diversity of foot-and-mouth disease serotype C virus in Kenya: evidence for probable vaccine strain re-introductions in the field. Epidemiology and infection 139, 189–196 (2011). URL http://www.ncbi.nlm.nih.gov/pubmed/20334728.

7. Thompson, D. et al. Economic costs of the foot and mouth disease outbreak in the United Kingdom in 2001. Revue Scientifique Et Technique De L Office International Des Epizooties 21, 675–687 (2002).

8. Perry, B. D. & Rich, K. M. Poverty impacts of foot-and-mouth disease and the poverty reduction implications of its control. Veterinary Record 160, 238–+ (2007).

9. Jemberu, W. T., Mourits, M., Rushton, J. & Hogeveen, H. Cost-benefit analysis of foot and mouth disease control in Ethiopia. Preventive Veterinary Medicine 132, 67–82 (2016). URL https://ac.els-cdn.com/S0167587716302896/1-s2.0-S0167587716302896-main.pdf?_tid=2c48a214-fc99-11e7-818f-00000aacb35f&acdnat=1516312226_72d77467f2afa7cb0ccd8e2e913dbf33.

10. Jemberu, W. T., Mourits, M. C. M., Woldehanna, T. & Hogeveen, H. Economic impact of foot and mouth disease outbreaks on smallholder farmers in Ethiopia. Preventive veterinary medicine 116, 26–36 (2014). URL http://www.ncbi.nlm.nih.gov/pubmed/24985154.

11. Lyons, N. A. et al. Impact of foot-and-mouth disease on mastitis and culling on a large-scale dairy farm in Kenya. Veterinary Research 46, 41 (2015). URL http://www.veterinaryresearch.org/content/46Z1/41.

12. Lyons, N. A. et al. Impact of foot-and-mouth disease on milk production on a large-scale dairy farm in Kenya. Preventive Veterinary Medicine 120, 177–186 (2015). URL http://dx.doi.org/10.1016/j.prevetmed.2015.04.004.

13. Qiu, Y. et al. Emergence of an exotic strain of serotype O foot-and-mouth disease virus O/ME-SA/Ind-2001d in South-East Asia in 2015. Transboundary and Emerging Diseases 65, e104–e112 (2018). URL http://doi.wiley.com/10.1111/tbed.12687.

14. Knowles, N. J. et al. Outbreaks of Foot-and-Mouth Disease in Libya and Saudi Arabia During 2013 Due to an Exotic 0/ME-SA/Ind-2001 Lineage Virus. Transboundary and Emerging Diseases 63, e431–e435 (2016). URL http://doi.wiley.com/10.1111/tbed.12299.

15. Roeder, P., Mariner, J. & Kock, R. Rinderpest: the veterinary perspective on eradication. Philosophical transactions of the Royal Society of London. Series B, Biological sciences 368, 20120139 (2013). URL http://www.ncbi.nlm.nih.goV/pubmed/23798687http://www.pubmedcentral.nih.gov/articlerender.fcgi?artid=PMC3720037.

16. Tekleghiorghis, T., Moormann, R. J. M., Weerdmeester, K. & Dekker, A. Foot-and-mouth Disease Transmission in Africa: Implications for Control, a Review. Transboundary and Emerging Diseases 1–16 (2014).

17. Hall, M. D., Knowles, N. J., Wadsworth, J., Rambaut, A. & Woolhouse, E. J. Reconstructing Geographical Movements and Host Species Transitions of Foot-and-Mouth Disease Virus Serotype SAT 2. mBio 4, 1–10 (2013).

18. Nardo, A. D., Knowles, N. J. & Paton, D. J. Combining livestock trade patterns with phylogenetics to help understand the spread of foot and mouth disease in sub-Saharan Africa, the Middle East and Southeast Asia. Revue Scientifique et Technique de l’Office International des Epizooties 30, 63–85 (2011).

19. Vosloo, W., Bastos, A. D. S., Sangare, O., Hargreaves, S. K. & Thomson, G. R. Review of the status and control of foot and mouth disease in sub-Saharan Africa. Revue Scientifique et Technique de l’Office International des Epizooties 21, 437–449 (2002).

20. Bronsvoort, B., Radford, A., Tanya, V. N., Kitching, R. P. & Morgan, K. L. The molecular epidemiology of foot-and-mouth disease viruses in the Adamawa Province of Cameroon. Journal of Clinical Microbiology 42, 2186–2196 (2004).

21. Habiela, M. et al. Molecular Characterization of Foot-and-Mouth Disease Viruses Collected from Sudan. Transboundary and Emerging Diseases 57, 305–314 (2010).

22. Sangare, O., Bastos, A. D. S., Venter, E. H. & Vosloo, W. Retrospective genetic analysis of SAT-1 type foot-and-mouth disease outbreaks in West Africa (1975-1981). Veterinary Microbiology 93, 279–289 (2003).

23. Couacy-Hymann, E. et al. Retrospective study of foot and mouth disease in West Africa from 1970 to 2003. Revue Scientifique Et Technique-Office International Des Epizooties 25, 1013–1024 (2006).

24. Jamal, S. M. & Belsham, G. J. Foot-and-mouth disease: past, present and future. Veterinary research 44, 116 (2013). URL http://www.ncbi.nlm.nih.gOv/pubmed/24308718http://www.pubmedcentral.nih.gov/articlerender.fcgi?artid=PMC4028749.

25. Reeve, R. et al. Sequence-Based Prediction for Vaccine Strain Selection and Identification of Antigenic Variability in Foot-and-Mouth Disease Virus. PLoS Computational Biology 6, e1001027 (2010).

26. Reeve, R. et al. Tracking the Antigenic Evolution of Foot-and-Mouth Disease Virus. PLOS ONE 11, e0159360 (2016). URL http://dx.plos.org/10.1371/journal.pone.0159360.

27. Motta, P. et al. Implications of the cattle trade network in Cameroon for regional disease prevention and control. Scientific Reports 7, 43932 (2017). URL http://www.nature.com/articles/srep43932.

28. Rady, A. A., Khalil, S. A. & Torky, H. A. Molecular Epidemiology of FMDV in Northern Egypt (2012-214). Alexandria Journal of Veterinary Sciences 41, 120–130 (2014). URL https://www.researchgate.net/profile/Helmy_Torky/publication/272669648_Molecular_Epidemiology_of_FMDV_in_Northern_Egypt_2012-214/links/58eeec11458515c4aa52d71e/Molecular-Epidemiology-of-FMDV-in-Northern-Egypt-2012-214.pdf.

29. Ferris, N. P. et al. Development and laboratory validation of a lateral flow device for the detection of foot-and-mouth disease virus in clinical samples. Journal of Virological Methods 155, 10–17 (2009).

30. Yang, M., Caterer, N. R., Xu, W. & Goolia, M. Development of a multiplex lateral flow strip test for foot-and-mouth disease virus detection using monoclonal antibodies. Journal of Virological Methods 221, 119–126 (2015). URL https://ac.els-cdn.com/S0166093415001779/1-s2.0-S0166093415001779-main.pdf?_tid=83f0b92a-fca3-11e7-8a38-00000aacb35f&acdnat=1516316668_3ab41f7d31aa131141069c9602420be8.

31. Fowler, V. L. et al. Recovery of Viral RNA and Infectious Foot-and-Mouth Disease Virus from Positive Lateral-Flow Devices (2014). URL http://eprints.gla.ac.uk/105850/.

32. Quick, J. et al. Real-time, portable genome sequencing for Ebola surveillance. Nature 530, 228–232 (2016). URL http://www.nature.com/articles/nature16996.

33. Ludi, A. et al. Serotype Diversity of Foot-and-Mouth-Disease Virus in Livestock without History of Vaccination in the Far North Region of Cameroon. Transboundary and Emerging Diseases 1–12 (2014).

34. Drummond, A. J., Suchard, M. A., Xie, D. & Rambaut, A. Bayesian phylogenetics with BEAUti and the BEAST 1.7. Molecular biology and evolution 29, 1969–73 (2012). URL http://www.ncbi.nlm.nih.gov/pubmed/22367748http://www.pubmedcentral.nih.gov/articlerender.fcgi?artid=PMC3408070.

35. Baele, G. et al. Improving the accuracy of demographic and molecular clock model comparison while accommodating phylogenetic uncertainty. Molecular biology and evolution 29, 2157–67 (2012). URL http://www.ncbi.nlm.nih.gov/pubmed/22403239http://www.pubmedcentral.nih.gov/articlerender.fcgi?artid=PMC3424409.

36. Baele, G., Li, W. L. S., Drummond, A. J., Suchard, M. A. & Lemey, P. Accurate model selection of relaxed molecular clocks in bayesian phylogenetics. Molecular biology and evolution 30, 239–43 (2013). URL http://www.ncbi.nlm.nih.gov/pubmed/23090976http://www.pubmedcentral.nih.gov/articlerender.fcgi?artid=PMC3548314.

37. Shapiro, B., Rambaut, A. & Drummond, A. J. Choosing Appropriate Substitution Models for the Phylogenetic Analysis of Protein-Coding Sequences. Molecular Biology and Evolution 23, 7–9 (2006). URL http://academic.oup.com/mbe/article/23/1/7/1193608/Choosing-Appropriate-Substitution-Models-for-the.

38. Gill, M.S. et al. Improving Bayesian population dynamics inference: a coalescent-based model for multiple loci. Molecular biology and evolution 30, 713–24 (2013). URL http://www.ncbi.nlm.nih.gov/pubmed/23180580http://www.pubmedcentral.nih.gov/articlerender.fcgi?artid=PMC3563973.

39. Lemey, P., Rambaut, A., Drummond, A. J. & Suchard, M. A. Bayesian Phylogeography Finds Its Roots. PLoS Computational Biology 5, e1000520 (2009). URL http://dx.plos.org/10.1371/journal.pcbi.1000520.

40. Lemey, P., Rambaut, A., Welch, J. J. & Suchard, M. A. Phylogeography takes a relaxed random walk in continuous space and time. Molecular biology and evolution 27, 1877–85 (2010). URL http://www.ncbi.nlm.nih.gov/pubmed/20203288 http://www.pubmedcentral.nih.gov/articlerender.fcgi?artid=PMC2915639.

41. Kosakovsky Pond, S. L. & Frost, S. D. W. Not So Different After All: A Comparison of Methods for Detecting Amino Acid Sites Under Selection. Molecular Biology and Evolution 22, 1208–1222 (2005). URL http://academic.oup.com/mbe/article/22/5/1208/1066893/Not-So-Different-After-All-A-Comparison-of-Methods.

42. Murrell, B. et al. FUBAR: A Fast, Unconstrained Bayesian AppRoximation for Inferring Selection. Molecular Biology and Evolution 30, 1196–1205 (2013). URL https://academic.oup.com/mbe/article-lookup/doi/10.1093/molbev/mst030.

43. Murrell, B. et al. Detecting individual sites subject to episodic diversifying selection. PLoS Genetics 8 (2012).

44. Kosakovsky Pond, S. L., Poon, A. F. Y., Leigh Brown, A. J. & Frost, S. D. W. A maximum likelihood method for detecting directional evolution in protein sequences and its application to influenza A virus. Molecular biology and evolution 25, 1809–24 (2008). URL http://www.ncbi.nlm.nih.gov/pubmed/18511426http://www.pubmedcentral.nih.gov/articlerender.fcgi?artid=PMC2515872.

45. Jones, D. T., Taylor, W. R. & Thornton, J. M. The rapid generation of mutation data matrices from protein sequences. Bioinformatics 8, 275–282 (1992). URL https://academic.oup.com/bioinformatics/article-lookup/doi/10.1093/bioinformatics/8.3.275.

46. Whelan, S. & Goldman, N. A General Empirical Model of Protein Evolution Derived from Multiple Protein Families Using a Maximum-Likelihood Approach. Molecular Biology and Evolution 18, 691–699 (2001). URL https://academic.oup.com/mbe/article-lookup/doi/10.1093/oxfordjournals.molbev.a003851.

