## Supplementary Material for "The evolution and phylodynamics of serotype A and SAT2 foot-and-mouth disease viruses in endemic regions of Africa"

#### ABSTRACT

Supplementary material containing additional details on the methods used for Field Work, Laboratory analysis, and supplementary figures S1-S8.

#### Methods

##### Field work

In June and July 2012 reported outbreak herds were visited in the Adamawa and the North West Regions of Cameroon. Using an established network of contacts in the local Ministry of Agriculture (MINEPIA) and veterinarian and para-veterinarians in the field herds were identified as having or recently having FMD (or believed to have had by the owners). All herds reported to the team during the field trip were visited by the team. A standard short questionnaire was used to record recent relevant history of the herd such as movements or purchases and permission was then sought to sample ~5 juvenile animals (<3 years old) animals per herd. The owner or herdsman was asked to select animals that he believed had lesions or had recently had signs consistent with FMD. Animals were cast in lateral recumbency and examined for clinical signs of FMD and an oropharngeal (probang) sample collected. Samples were suspended in 20ml of buffered phosphate and 1.8 ml of fluid was then drawn off with a Pasteur pipette trying to ensure plenty of cellular material and decanted to a 1.8ml cryovial and immediately stored down in a liquid nitrogen shipper[1,2].

This work was carried out with Cameroonian Ministère de l'Élevage des Pêches et Industries Animales (MINEPIA) staff, following local guidelines and regulations, as part of their disease surveillance activities and had approval from the Cameroonian Academy of Science.

##### Laboratory analysis of isolates collected in 2012

Virus isolation was attempted from all samples in either primary bovine thyroid cell cultures for all epithelial and oropharyngeal samples or renal swine cell cultures (RS) for epithelial and pig whole-blood samples by the WRL standard protocol[3]. All cultures showing FMDV cytopathic effect were harvested and serotyped with the WRL indirect sandwich enzyme-linked immunosorbent assay[3–5].

All viruses that were sequenced underwent a maximum of two passages in cell culture. The viral RNA was extracted from 460 $\mu$l of either the original viral suspension or the tissue culture supernatant, following the manufacturer's instructions (RNeasy Protect minikit; Qiagen Ltd.), eluted into 50 $\mu$l of diethyl pyrocarbonate-treated $H_2O$, and stored at $-70\,^{\circ}$C until used for reverse transcription. A one-stage reverse transcription-PCR was used to amplify the 3' end of the VP1 gene from 5 $\mu$l of purified RNA according to the manufacturer's instructions (Ready-To-Go reverse transcription-PCR tube; Amersham Pharmacia Biotech United Kingdom Ltd.). The primers were used at a concentration of 0.2 to 0.5 pmol/$\mu$l and depended on the FMDV serotype[2]. For the reverse transcription phase, the samples were incubated at 42 °C for 30 min, followed by 5 min at 90 °C to inactivate the reverse transcriptase. After further incubation at 94 °C for 3 min, the samples were subjected to 30 cycles of PCR, with 1 cycle consisting of template denaturation at 94 °C for 1 min, primer annealing at 55 °C for 1 min, and primer extension at 72 °C for 1.5 min. There was a final incubation at 72 °C for 5 min. Amplicons were purified to remove unincorporated primers and nucleotides (Wizard Prep DNA purification kit; Promega) and extracted with phenol-chloroform to remove any viral protein or infectivity. The purified amplicons were sequenced by MWG-Biotech as recommended by the manufacturer (ABI Prism BigDye Terminators version 3.0 cycle sequencing kits; Applied Biosystems). All amplicons were forward sequenced with the PCR primer for that serotype and reverse sequenced with internal primer NK72.

A total of 19 herds were reported to the team during June/July 2012 (9 from the Adamawa and 10 from the North West Regions), and from these 87 animals were clinically examined and probang samples collected. Virus was successfully cultured and sequenced from 18/87 samples, 9 SAT2 positive animals and 9 A positive animals. The 9 animals from which SAT2 was recovered had foot lesions and 8/9 also had mouth lesions. Similarly 8/9 animals from which serotype A was recovered had foot lesions and 7/9 had mouth lesions. Of those for which no virus was cultured 59/69 had foot lesions and 52/69 had mouth lesions. The sample codes and accession numbers of the sequenced samples are given in Table 1 in the main paper.

In order to relate the new sequences to previously circulating strains, further A and SAT2 VP1 sequences were accessed from our previous study[2] and combined with other publicly available sequences from Genbank (http://www.ncbi.nlm.nih.gov/genbank/) as well as recent sequences provided by Ludi et al.[6]. In total there were sequences available for Cameroon from 2000, 2005, 2012, 2013 and 2014.

#### References


1. Kitching, R. P. & Donaldson, A. I. Collection and transportation of specimens for vesicular virus investigation. *Revue Sci. et Tech. de l'Office Int. des Epizoot.* **6**, 263–272 (1987).
2. Bronsvoort, B., Radford, A., Tanya, V. N., Kitching, R. P. & Morgan, K. L. The molecular epidemiology of foot-and-mouth disease viruses in the Adamawa Province of Cameroon. *J. Clin. Microbiol.* **42**, 2186–2196 (2004).
3. Anon. Chapter 2.1.1, Foot and Mouth Disease. In *Manual of Diagnostic Tests and Vaccines for Terrestrial Animals*, vol. 2005 (O.I.E., 2005), 5th edn.
4. Ferris, N. P., Powell, H. & Donaldson, A. I. Use of Pre-Coated Immunoplates and Freeze-Dried Reagents For the Diagnosis of Foot-and-Mouth-Disease and Swine Vesicular Disease By Enzyme-Linked Immunosorbent-Assay (Elisa). *J. Virol. Methods* **19**, 197–206 (1988).
5. Roeder, P. L. & Le Blanc Smith, P. M. Detection and typing of foot-and-mouth disease virus by enzyme-linked immunosorbent assay: a sensitive, rapid and reliable technique for primary diagnosis. *Res. veterinary science* **43**, 225–32 (1987).
6. Ludi, A. *et al.* Serotype Diversity of Foot-and-Mouth-Disease Virus in Livestock without History of Vaccination in the Far North Region of Cameroon. *Transboundary Emerg. Dis.* 1–12 (2014). DOI 10.1111/tbed.12227.

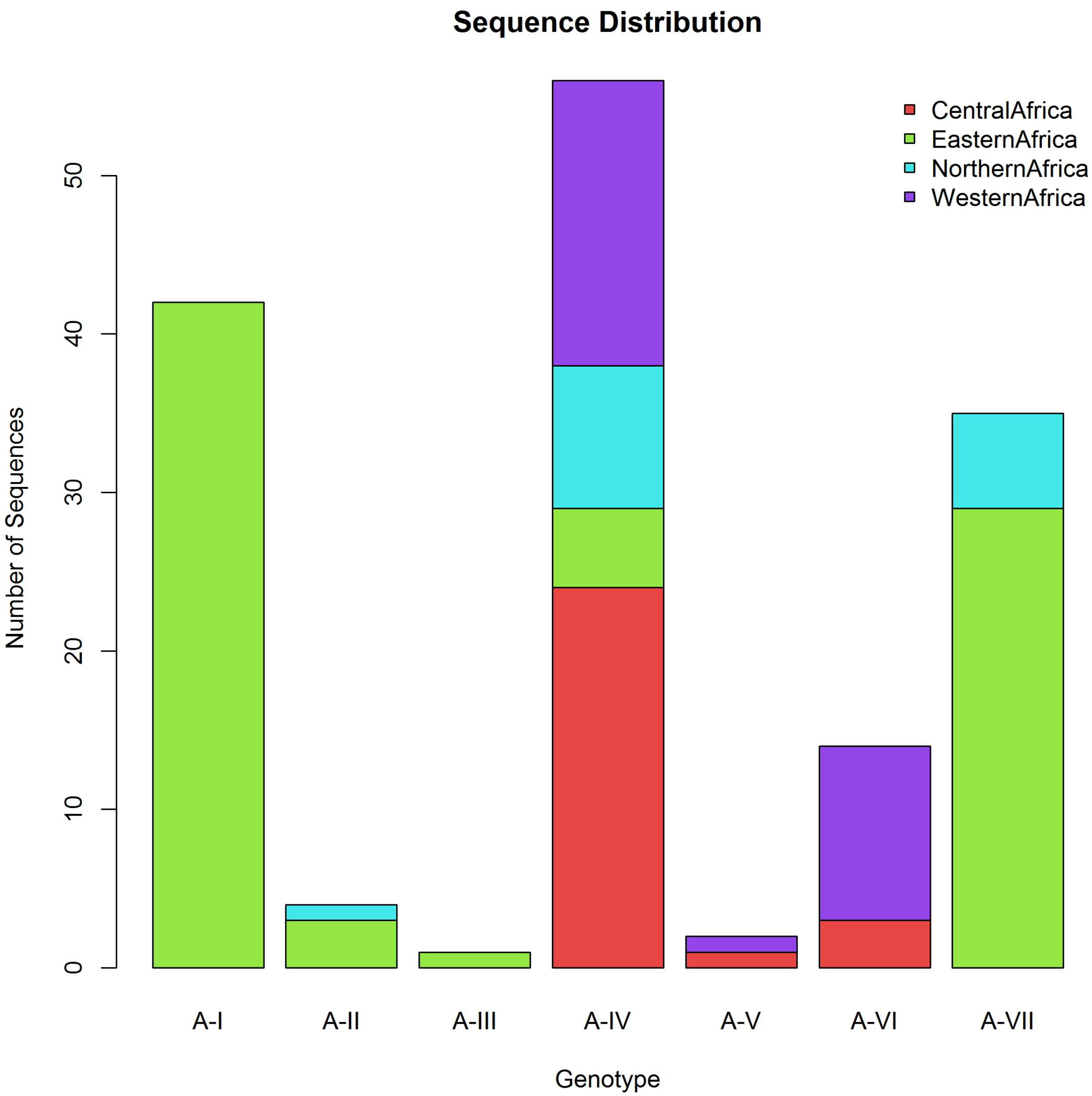


**Figure S1.** Composition of 154 serotype A sequence dataset by genotype and region.

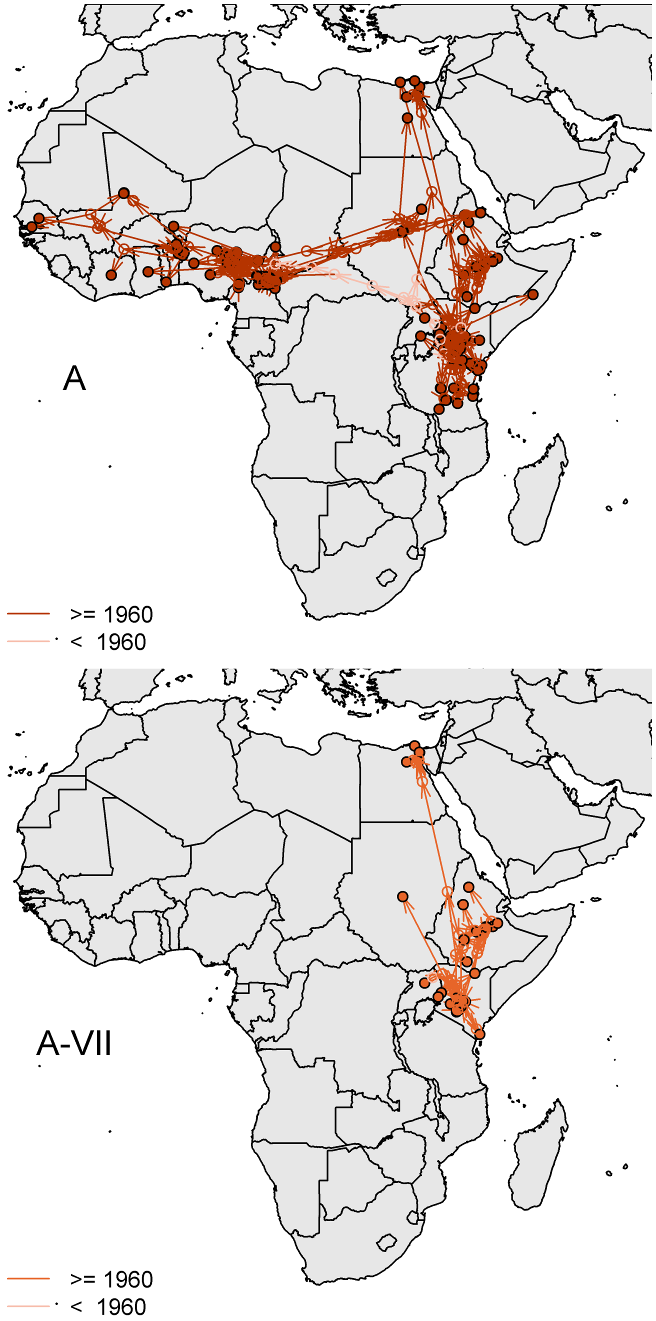


**Figure S2. Spatial projection of maximum clade credibility trees from serotype A sequences.** Spatial components were inferred using homogenous Brownian motion and projected onto maps. Upper left: MCC tree from all serotype A dataset. Upper right: MCC tree from A-I sequences only, Lower left: A-VII, Lower right: A-IV. Filled circles are the positions of the sequences, open circles are the inferred positions of the ancestral nodes. Dark arrows indicate tree branches and inferred transmission routes with dates $\geq$ 1960, pale arrows indicate older routes (< 1960).

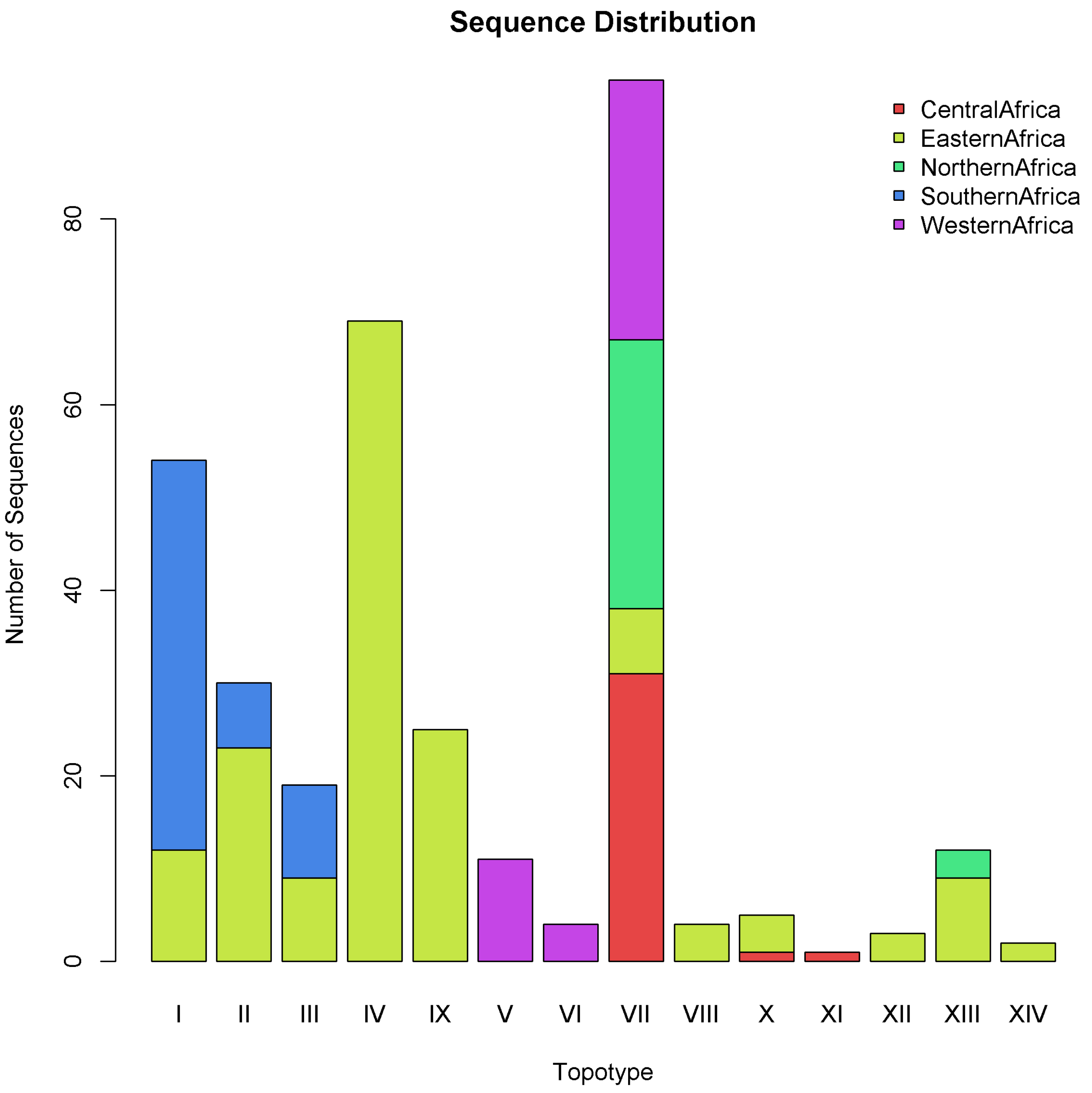


**Figure S3.** Composition of 334 serotype SAT2 sequence data by topotype and region.

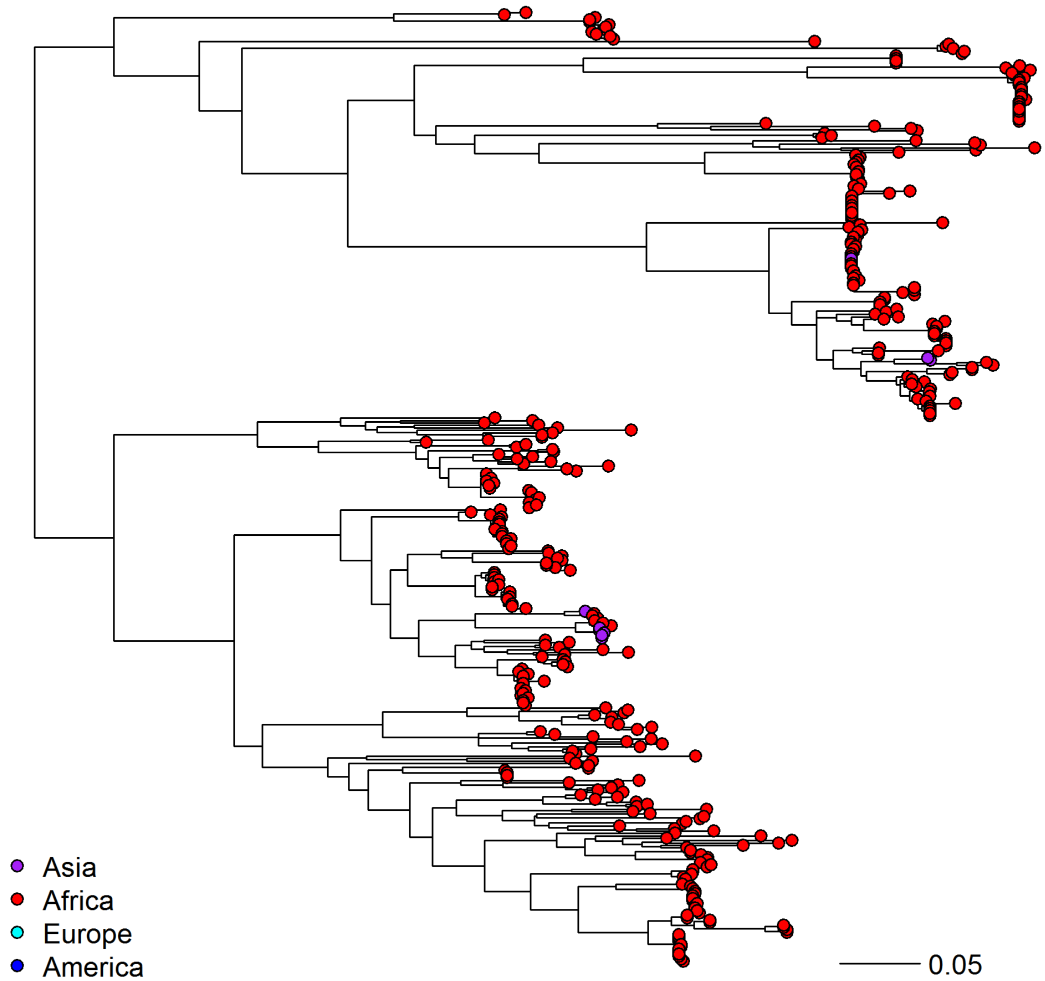


**Figure S4. Maximum likelihood tree from world wide SAT2 sequences**, showing the majority of sequences are from Africa, with only occasional incursions into Asia.

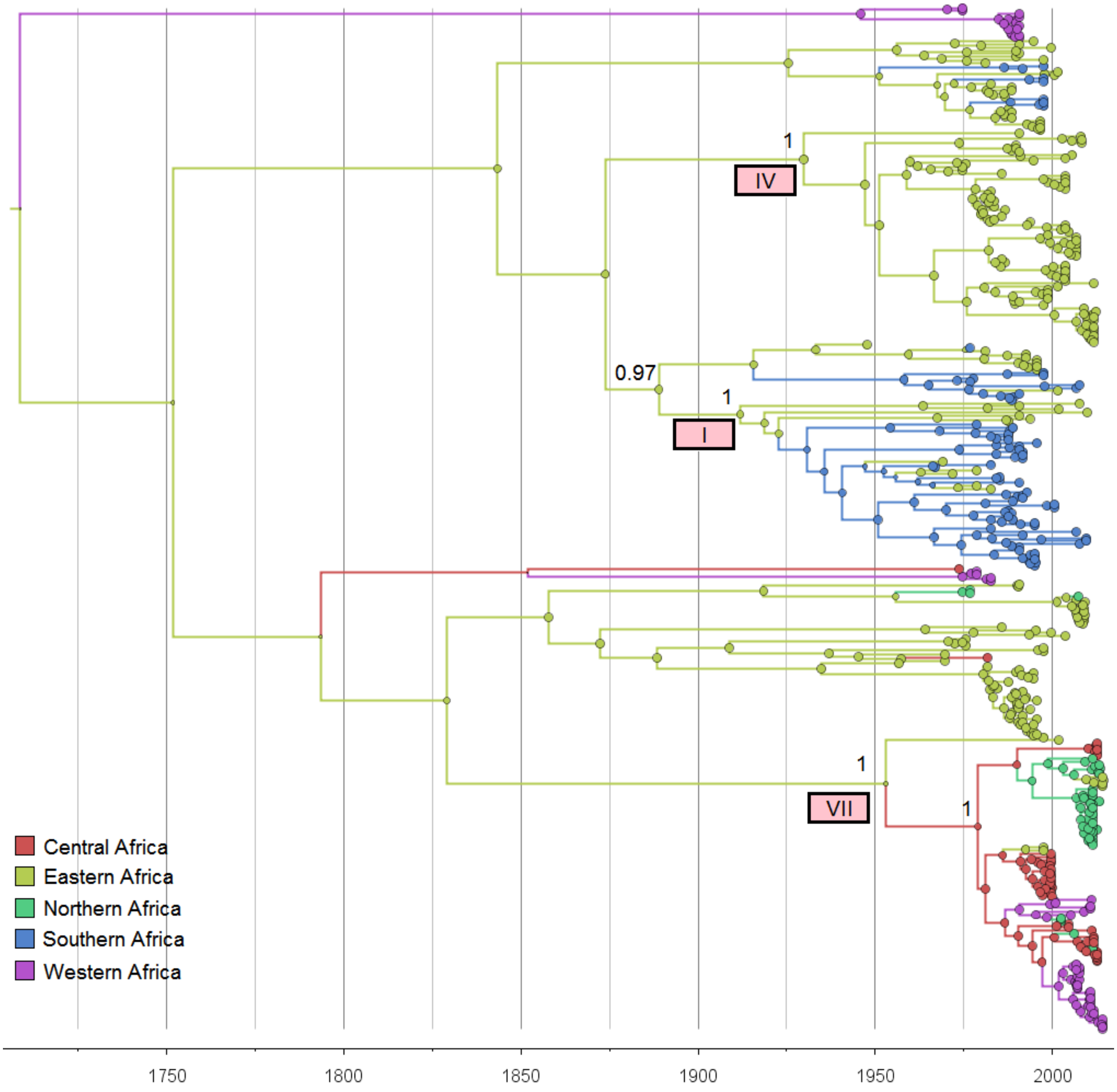


**Figure S5. Time scaled tree of 334 SAT2 sequences from Africa**, with genotypes used in further analysis marked.

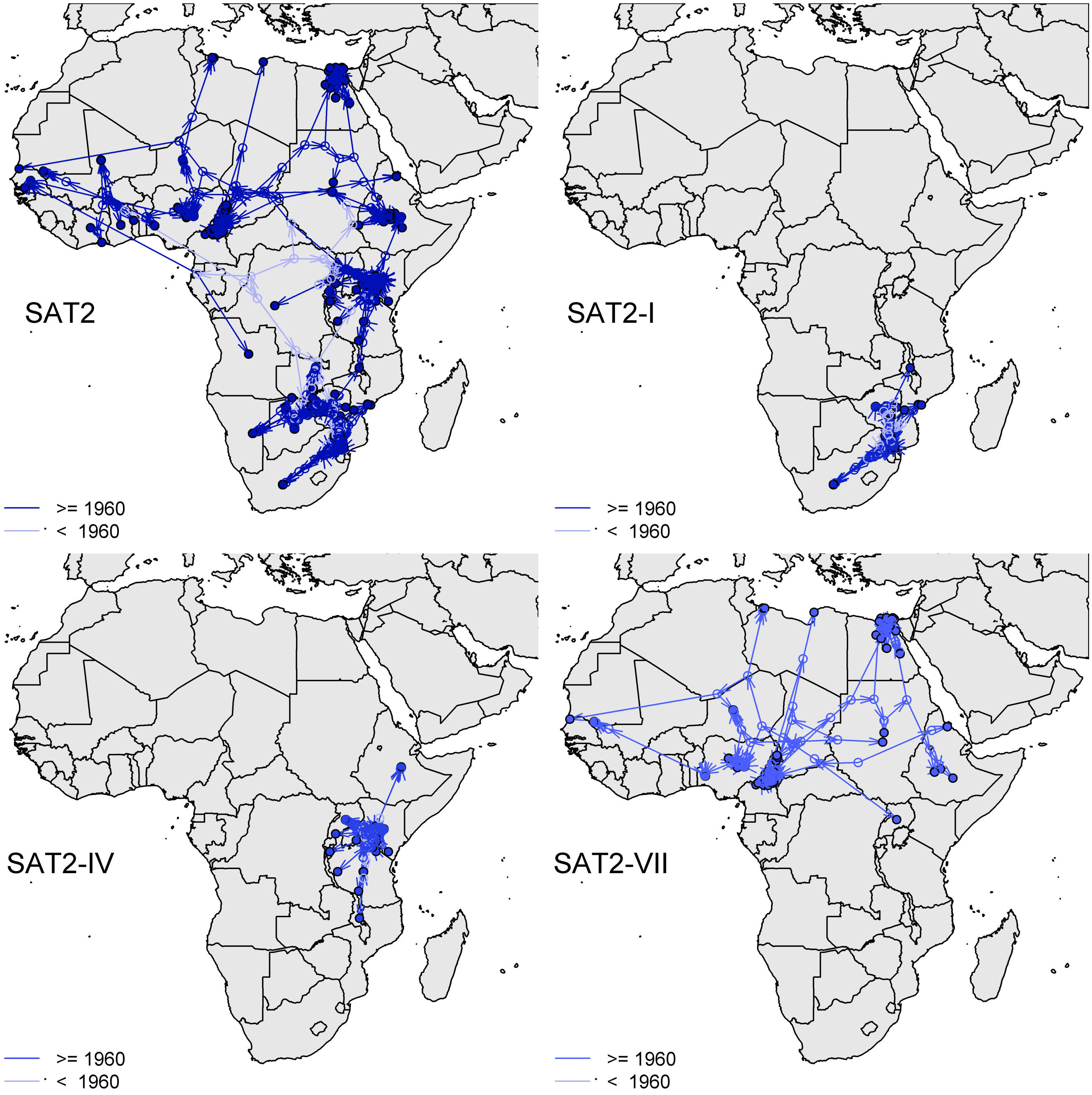


**Figure S6. Spatial projection of maximum clade credibility trees from serotype SAT2 sequences.** Spatial components were inferred using homogenous Brownian motion and projected onto maps. Upper left: MCC tree from all serotype SAT2 dataset. Upper right: MCC tree from Topotype SAT2-I sequences only, Lower left: SAT2-IV, Lower right: SAT2-VII. Filled circles are the positions of the sequences, open circles are the inferred positions of the ancestral nodes. Dark arrows indicate tree branches and inferred transmission routes with dates $\geq$ 1960, pale arrows indicate older routes ($<$ 1960).

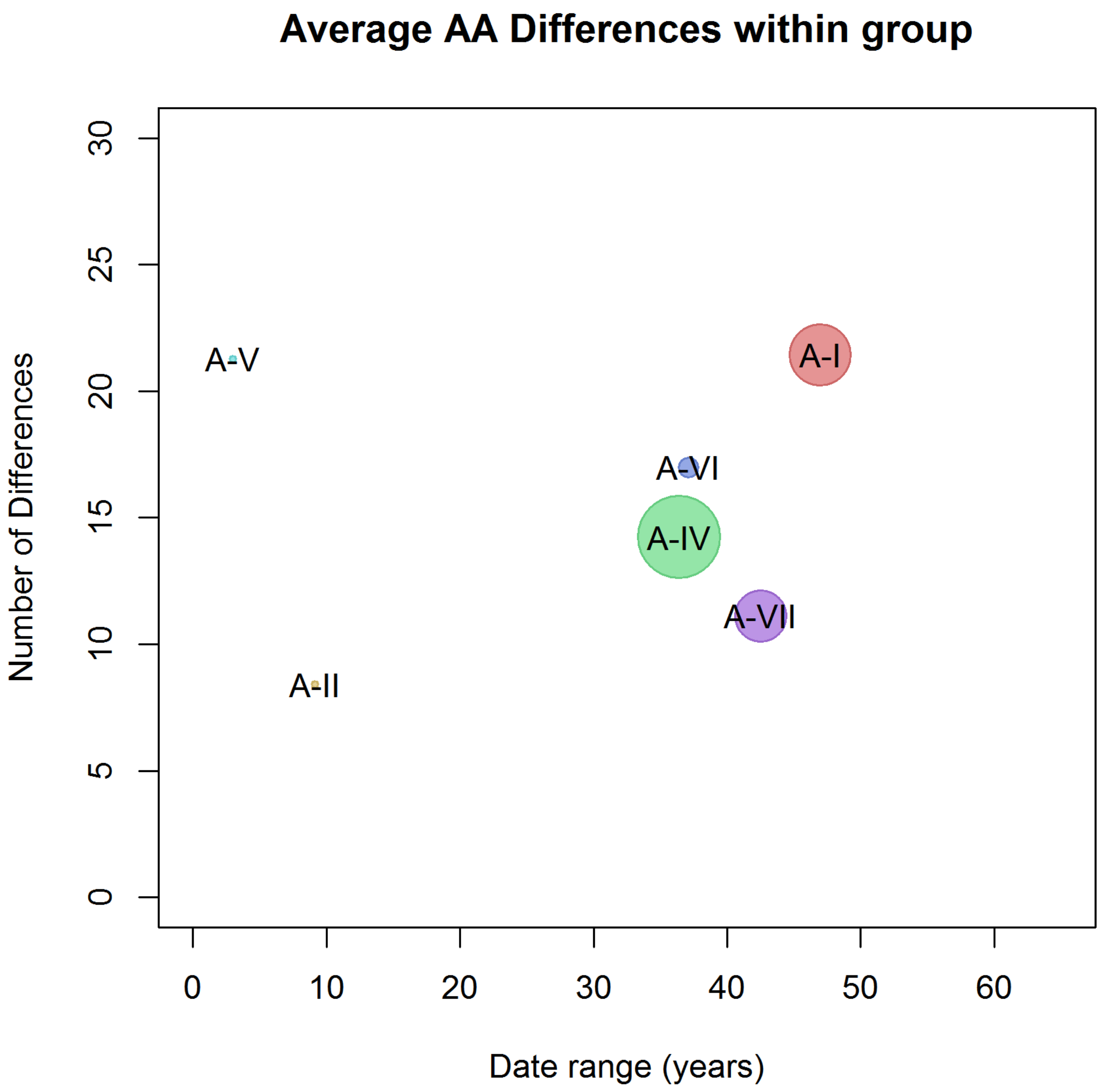


**Figure S7.** Average number of pairwise amino acid differences for serotype A within A groups as a function of the date range of the sequences (youngest – oldest). Circle size indicates the number of sequences within each group (see Figure S1).

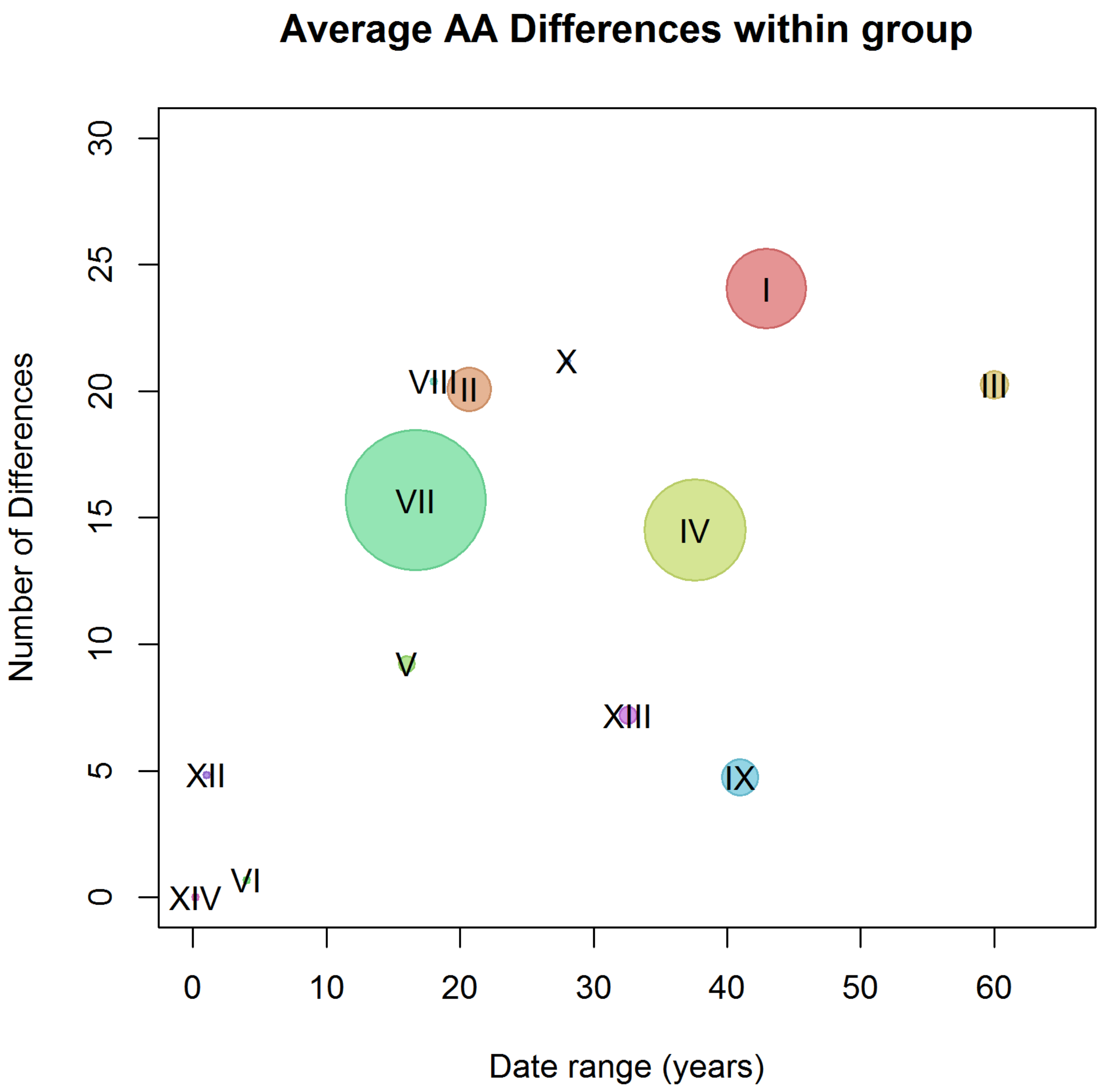


**Figure S8.** Average number of pairwise amino acid differences for serotype SAT2 within Topotypes as a function of the date range of the sequences (youngest – oldest). Circle size indicates the number of sequences within each group (see Figure S2).

#### A and SAT2 within and between group amino acid variations

Within group stats for A

| Group (A) | NumSeqs | Mean | Median | StDev | MinDate | MaxDate |
|---|---|---|---|---|---|---|
| (ALL) | 154 | 0.112 | 0.118 | 0.034 | 1964.500 | 2013.888 |
| A-I | 42 | 0.096 | 0.103 | 0.040 | 1966.500 | 2013.500 |
| A-II | 4 | 0.038 | 0.038 | 0.034 | 1972.362 | 1981.500 |
| A-III | 1 | NA | NA | NA | 1964.500 | 1964.500 |
| A-IV | 56 | 0.063 | 0.066 | 0.023 | 1977.500 | 2013.888 |
| A-V | 2 | 0.095 | 0.095 | NA | 1973.500 | 1976.500 |
| A-VI | 14 | 0.076 | 0.085 | 0.032 | 1973.500 | 2010.608 |
| A-VII | 35 | 0.050 | 0.052 | 0.017 | 1966.096 | 2008.581 |

Within Topotype Stats for SAT2

| TopoType | NumSeqs | Mean | Median | StDev | MinDate | MaxDate |
|---|---|---|---|---|---|---|
| (ALL) | 334 | 0.162 | 0.176 | 0.049 | 1948.500 | 2015.189 |
| I | 54 | 0.107 | 0.108 | 0.028 | 1967.770 | 2010.701 |
| II | 30 | 0.090 | 0.094 | 0.041 | 1981.847 | 2002.500 |
| III | 19 | 0.090 | 0.097 | 0.033 | 1948.500 | 2008.500 |
| IV | 69 | 0.065 | 0.069 | 0.024 | 1975.500 | 2013.104 |
| V | 11 | 0.041 | 0.028 | 0.034 | 1975.500 | 1991.500 |
| VI | 4 | 0.003 | 0.005 | 0.002 | 1979.500 | 1983.500 |
| VII | 95 | 0.070 | 0.074 | 0.029 | 1998.500 | 2015.189 |
| VIII | 4 | 0.091 | 0.097 | 0.027 | 1986.500 | 2004.500 |
| IX | 25 | 0.021 | 0.005 | 0.035 | 1957.500 | 1998.500 |
| X | 5 | 0.095 | 0.102 | 0.029 | 1970.500 | 1998.500 |
| XI | 1 | NA | NA | NA | 1974.500 | 1974.500 |
| XII | 3 | 0.022 | 0.023 | 0.003 | 1975.500 | 1976.500 |
| XIII | 12 | 0.032 | 0.009 | 0.040 | 1977.500 | 2010.066 |
| XIV | 2 | 0.000 | 0.000 | NA | 1991.329 | 1991.500 |

Between Group Stats for A

| Group | A-I | A-II | A-III | A-IV | A-V | A-VI | A-VII |
|---|---|---|---|---|---|---|---|
| A-I | 0.096 | 0.135 | 0.122 | 0.130 | 0.121 | 0.121 | 0.128 |
| A-II | 0.135 | 0.038 | 0.123 | 0.134 | 0.118 | 0.125 | 0.130 |
| A-III | 0.122 | 0.123 | NA | 0.114 | 0.100 | 0.120 | 0.116 |
| A-IV | 0.130 | 0.134 | 0.114 | 0.063 | 0.114 | 0.121 | 0.131 |
| A-V | 0.121 | 0.118 | 0.100 | 0.114 | 0.095 | 0.097 | 0.113 |
| A-VI | 0.121 | 0.125 | 0.120 | 0.121 | 0.097 | 0.076 | 0.107 |
| A-VII | 0.128 | 0.130 | 0.116 | 0.131 | 0.113 | 0.107 | 0.050 |

Between Topotype Stats for SAT2

| TopoType | I | II | III | IV | V | VI | VII | VIII | IX | X | XI | XII | XIII | XIV |
|---|---|---|---|---|---|---|---|---|---|---|---|---|---|---|
| I | 0.107 | 0.167 | 0.143 | 0.141 | 0.152 | 0.179 | 0.186 | 0.168 | 0.180 | 0.179 | 0.217 | 0.187 | 0.186 | 0.163 |
| II | 0.167 | 0.090 | 0.167 | 0.176 | 0.178 | 0.196 | 0.223 | 0.200 | 0.195 | 0.213 | 0.201 | 0.220 | 0.205 | 0.192 |
| III | 0.143 | 0.167 | 0.090 | 0.135 | 0.163 | 0.188 | 0.199 | 0.183 | 0.165 | 0.193 | 0.212 | 0.207 | 0.191 | 0.170 |
| IV | 0.141 | 0.176 | 0.135 | 0.065 | 0.173 | 0.191 | 0.188 | 0.172 | 0.185 | 0.192 | 0.212 | 0.201 | 0.190 | 0.178 |
| V | 0.152 | 0.178 | 0.163 | 0.173 | 0.041 | 0.166 | 0.186 | 0.142 | 0.145 | 0.159 | 0.208 | 0.169 | 0.165 | 0.151 |
| VI | 0.179 | 0.196 | 0.188 | 0.191 | 0.166 | 0.003 | 0.234 | 0.208 | 0.204 | 0.210 | 0.222 | 0.225 | 0.224 | 0.201 |
| VII | 0.186 | 0.223 | 0.199 | 0.188 | 0.186 | 0.234 | 0.070 | 0.171 | 0.181 | 0.177 | 0.237 | 0.201 | 0.177 | 0.174 |
| VIII | 0.168 | 0.200 | 0.183 | 0.172 | 0.142 | 0.208 | 0.171 | 0.091 | 0.141 | 0.139 | 0.218 | 0.142 | 0.151 | 0.159 |
| IX | 0.180 | 0.195 | 0.165 | 0.185 | 0.145 | 0.204 | 0.181 | 0.141 | 0.021 | 0.128 | 0.217 | 0.154 | 0.163 | 0.127 |
| X | 0.179 | 0.213 | 0.193 | 0.192 | 0.159 | 0.210 | 0.177 | 0.139 | 0.128 | 0.095 | 0.235 | 0.143 | 0.175 | 0.147 |
| XI | 0.217 | 0.201 | 0.212 | 0.212 | 0.208 | 0.222 | 0.237 | 0.218 | 0.217 | 0.235 | NA | 0.247 | 0.221 | 0.222 |
| XII | 0.187 | 0.220 | 0.207 | 0.201 | 0.169 | 0.225 | 0.201 | 0.142 | 0.154 | 0.143 | 0.247 | 0.022 | 0.188 | 0.165 |
| XIII | 0.186 | 0.205 | 0.191 | 0.190 | 0.165 | 0.224 | 0.177 | 0.151 | 0.163 | 0.175 | 0.221 | 0.188 | 0.032 | 0.149 |
| XIV | 0.163 | 0.192 | 0.170 | 0.178 | 0.151 | 0.201 | 0.174 | 0.159 | 0.127 | 0.147 | 0.222 | 0.165 | 0.149 | 0.000 |

No. Sites 223

Between groups in number of AA changes for A

| Group | A-I | A-II | A-III | A-IV | A-V | A-VI | A-VII |
|---|---|---|---|---|---|---|---|
| A-I | 21 | 30 | 27 | 29 | 27 | 27 | 28 |
| A-II | 30 | 8 | 27 | 30 | 26 | 28 | 29 |
| A-III | 27 | 27 | NA | 25 | 22 | 27 | 26 |
| A-IV | 29 | 30 | 25 | 14 | 25 | 27 | 29 |
| A-V | 27 | 26 | 22 | 25 | 21 | 22 | 25 |
| A-VI | 27 | 28 | 27 | 27 | 22 | 17 | 24 |
| A-VII | 28 | 29 | 26 | 29 | 25 | 24 | 11 |

No. Sites 224

Between topotype in number of AA changes for SAT2

| TopoType | I | II | III | IV | V | VI | VII | VIII | IX | X | XI | XII | XIII | XIV |
|---|---|---|---|---|---|---|---|---|---|---|---|---|---|---|
| I | 24 | 37 | 32 | 32 | 34 | 40 | 42 | 38 | 40 | 40 | 49 | 42 | 42 | 36 |
| II | 37 | 20 | 37 | 39 | 40 | 44 | 50 | 45 | 44 | 48 | 45 | 49 | 46 | 43 |
| III | 32 | 37 | 20 | 30 | 36 | 42 | 45 | 41 | 37 | 43 | 47 | 46 | 43 | 38 |
| IV | 32 | 39 | 30 | 15 | 39 | 43 | 42 | 39 | 41 | 43 | 48 | 45 | 43 | 40 |
| V | 34 | 40 | 36 | 39 | 9 | 37 | 42 | 32 | 33 | 36 | 46 | 38 | 37 | 34 |
| VI | 40 | 44 | 42 | 43 | 37 | 1 | 52 | 47 | 46 | 47 | 50 | 50 | 50 | 45 |
| VII | 42 | 50 | 45 | 42 | 42 | 52 | 16 | 38 | 41 | 40 | 53 | 45 | 40 | 39 |
| VIII | 38 | 45 | 41 | 39 | 32 | 47 | 38 | 20 | 32 | 31 | 49 | 32 | 34 | 36 |
| IX | 40 | 44 | 37 | 41 | 33 | 46 | 41 | 32 | 5 | 29 | 49 | 34 | 36 | 29 |
| X | 40 | 48 | 43 | 43 | 36 | 47 | 40 | 31 | 29 | 21 | 53 | 32 | 39 | 33 |
| XI | 49 | 45 | 47 | 48 | 46 | 50 | 53 | 49 | 49 | 53 | NA | 55 | 50 | 50 |
| XII | 42 | 49 | 46 | 45 | 38 | 50 | 45 | 32 | 34 | 32 | 55 | 5 | 42 | 37 |
| XIII | 42 | 46 | 43 | 43 | 37 | 50 | 40 | 34 | 36 | 39 | 50 | 42 | 7 | 33 |
| XIV | 36 | 43 | 38 | 40 | 34 | 45 | 39 | 36 | 29 | 33 | 50 | 37 | 33 | 0 |

**Figure S9.** Table of within and between group amino acid differences for A and SAT2

### Comparison of A and SAT2 clade diversities

| **Clock Rate** | **A** | **A-I** | **A-VII** | **A-IV** | **SAT2** | **SAT2-IV** | **SAT2-I** | **SAT2-VII** |
|---|---|---|---|---|---|---|---|---|
| Mean | 0.00566 | 0.00431 | 0.00401 | 0.00865 | 0.00320 | 0.00066 | 0.00302 | 0.00743 |
| Median | 0.00563 | 0.00407 | 0.00387 | 0.00857 | 0.00318 | 0.00065 | 0.00291 | 0.00727 |
| Q1 | 0.00519 | 0.00321 | 0.00321 | 0.00805 | 0.00289 | 0.00058 | 0.00232 | 0.00651 |
| Q3 | 0.00610 | 0.00517 | 0.00459 | 0.00919 | 0.00348 | 0.00072 | 0.00357 | 0.00820 |
| Lower | 0.00408 | 0.00141 | 0.00153 | 0.00638 | 0.00206 | 0.00037 | 0.00104 | 0.00443 |
| Upper | 0.00744 | 0.00793 | 0.00663 | 0.01090 | 0.00437 | 0.00093 | 0.00542 | 0.01071 |
| Lower 95% HPD | 0.00447 | 0.00159 | 0.00223 | 0.00711 | 0.00240 | 0.00045 | 0.00148 | 0.00501 |
| Upper 95% HPD | 0.00685 | 0.00736 | 0.00642 | 0.01053 | 0.00407 | 0.00090 | 0.00475 | 0.01013 |

| **Root Age** | **A** | **A-I** | **A-VII** | **A-IV** | **SAT2** | **SAT2-IV** | **SAT2-I** | **SAT2-VII** |
|---|---|---|---|---|---|---|---|---|
| Mean | 1936.4 | 1921.6 | 1955.1 | 1974.9 | 1693.0 | 1752.0 | 1900.0 | 1977.3 |
| Median | 1937.3 | 1926.4 | 1956.7 | 1974.9 | 1709.1 | 1759.9 | 1905.2 | 1982.7 |
| Q1 | 1932.6 | 1908.8 | 1951.5 | 1974.0 | 1655.3 | 1723.2 | 1883.5 | 1972.8 |
| Q3 | 1941.5 | 1939.3 | 1960.4 | 1975.8 | 1752.2 | 1790.1 | 1919.0 | 1988.5 |
| Lower | 1919.2 | 1863.0 | 1938.2 | 1971.5 | 1510.1 | 1624.1 | 1830.3 | 1949.3 |
| Upper | 1953.6 | 1960.7 | 1966.1 | 1977.5 | 1867.0 | 1881.1 | 1952.9 | 1995.7 |
| Lower 95% HPD | 1922.1 | 1875.2 | 1940.0 | 1972.7 | 1496.8 | 1626.9 | 1849.1 | 1944.4 |
| Upper 95% HPD | 1949.7 | 1955.5 | 1966.1 | 1977.5 | 1810.3 | 1846.3 | 1943.2 | 1995.1 |

| **Number of AA Differences** | **A** | **A-I** | **A-VII** | **A-IV** | **SAT2** | **SAT2-IV** | **SAT2-I** | **SAT2-VII** |
|---|---|---|---|---|---|---|---|---|
| Mean | 25.0 | 21.4 | 11.1 | 14.2 | 36.3 | 14.5 | 24.0 | 15.7 |
| Median | 26.4 | 23.1 | 11.6 | 14.7 | 39.4 | 15.6 | 24.2 | 16.6 |
| Q1 | 22.1 | 17.8 | 9.5 | 11.3 | 32.1 | 12.4 | 20.5 | 12.4 |
| Q3 | 30.5 | 28.4 | 13.7 | 17.9 | 43.6 | 17.7 | 28.0 | 19.0 |
| Lower | 9.5 | 2.1 | 3.2 | 2.1 | 15.1 | 5.2 | 9.4 | 3.1 |
| Upper | 41.3 | 37.9 | 20.0 | 25.0 | 59.7 | 25.4 | 39.0 | 28.7 |
| Lower 95% HPD | 8.4 | 2.1 | 3.1 | 5.0 | 13.1 | 1.0 | 11.4 | 1.0 |
| Upper 95% HPD | 36.3 | 35.8 | 18.0 | 23.8 | 54.4 | 22.1 | 35.9 | 28.0 |

| **Diffusion rate** | **A** | **A-I** | **A-VII** | **A-IV** | **SAT2** | **SAT2-IV** | **SAT2-I** | **SAT2-VII** |
|---|---|---|---|---|---|---|---|---|
| Mean | 101.8 | 33.7 | 110.4 | 209.0 | 46.1 | 8.8 | 20.3 | 327.6 |
| Median | 101.3 | 32.7 | 108.3 | 205.9 | 45.2 | 8.6 | 19.4 | 322.8 |
| Q1 | 94.7 | 25.8 | 95.0 | 190.5 | 41.0 | 7.7 | 16.0 | 286.0 |
| Q3 | 108.7 | 41.3 | 125.3 | 221.7 | 49.8 | 9.6 | 23.8 | 371.8 |
| Lower | 73.8 | 9.8 | 51.3 | 146.8 | 28.3 | 5.0 | 6.5 | 160.3 |
| Upper | 129.4 | 63.7 | 167.0 | 267.0 | 62.7 | 12.5 | 35.2 | 500.1 |
| Lower 95% HPD | 81.1 | 14.7 | 72.2 | 166.1 | 32.5 | 6.0 | 10.3 | 200.7 |
| Upper 95% HPD | 123.2 | 52.4 | 153.6 | 260.7 | 60.2 | 12.0 | 32.7 | 451.1 |



**Figure S10.** Comparison of clock rate, root age (TMRCA), number of amino acid differences and diffusion rate for A and

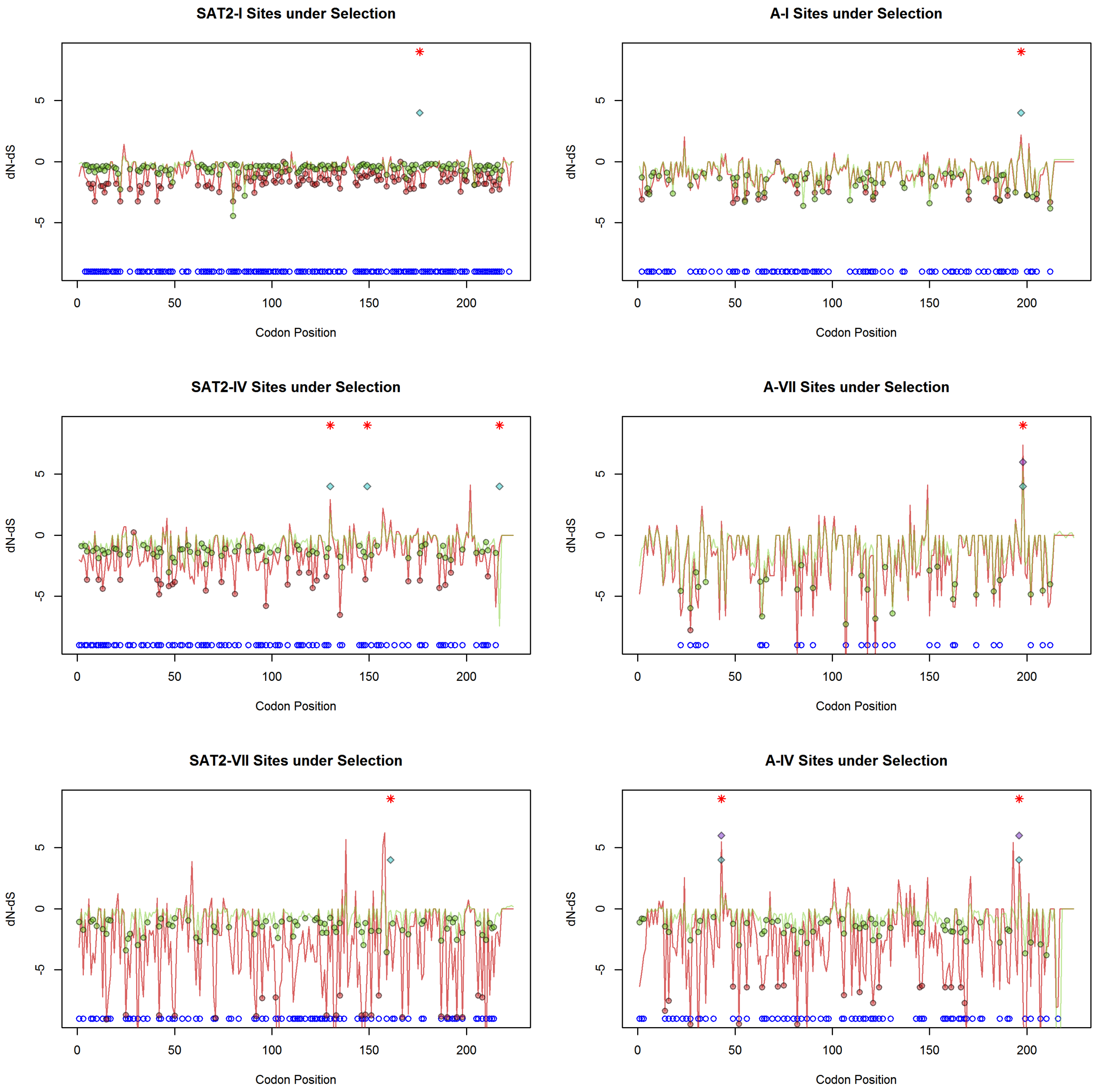


**Figure S11.** Sites under positive and negative selection from SLAC (red), FEL (green), MEME (blue) and FUBAR (purple). The normalised dN-dS values for SLAC and FEL are plotted, and significantly selected sites (p-value $\leq 0.01$) are indicated with a circle. The sites under selection using MEME (p-value $\leq 0.01$) and FUBAR sites (probability $\geq 0.99$) are indicated with blue and purple diamonds respectively. Positively selected sites, significant under at least one test are indicated with a red star, and significant negatively selected sites are indicated with a blue circle.
